# Principles of sensor-effector organization in six-transmembrane ion channels

**DOI:** 10.1101/2021.08.03.454958

**Authors:** Alex Dou, Po Wei Kang, Panpan Hou, Mark A. Zaydman, Jie Zheng, Timothy Jegla, Jianmin Cui

## Abstract

Receptor proteins sense stimuli and generate downstream signals via sensor and effector domains. Presently, the structural constraints on sensor-effector organization across receptor protein superfamilies are not clear. Here, we perform statistical coupling analysis (SCA) on the transient receptor potential (TRP) and voltage-gated potassium (Kv) ion channel superfamilies to characterize the networks of coevolving residues, or protein sectors, that mediate their receptor functions. Comparisons to structural and functional studies reveal a conserved “core” sector that extends from the pore and mediates effector functions, including pore gating and sensor-pore coupling, while sensors correspond to family-specific “accessory” sectors and localize according to three principles: Sensors (1) may emerge in any region with access to the core, (2) must maintain contact with the core, and (3) must preserve the integrity of the core. This sensor-core architecture may represent a conserved and generalizable paradigm for the structure-function relationships underlying the evolution of receptor proteins.

## Introduction

Receptor proteins transduce physiologically relevant stimuli into intracellular signals. In these molecules, signal transduction proceeds through three steps (Fig. 1A). First, a stimulus activates a sensor domain. Second, the activation of the sensor is directly or allosterically propagated to an effector domain. Finally, the effector initiates a signal that produces a downstream cellular response. Although these processes have been studied in individual receptor families, general principles describing the organization of sensors and effectors across receptor superfamilies have yet to be elucidated.

**Figure 1.**
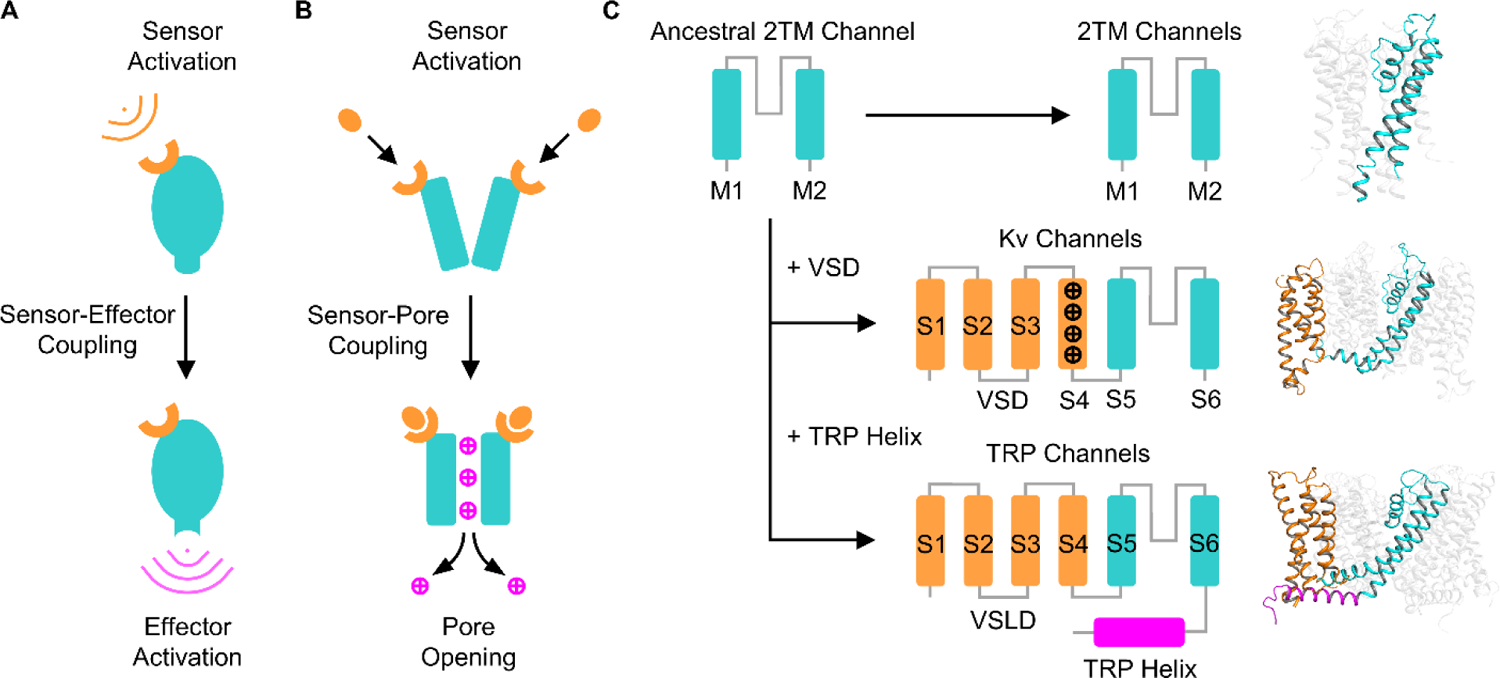
Six-transmembrane channel gating and evolution. (A) Schematic representation of signal transduction in receptor proteins. (B) Signal transduction in gated ion channels. (C) Evolution of six-transmembrane (6TM) channel structure. The addition of four transmembrane helices (S1-S4) to the N-terminus of an ancestral two-transmembrane (2TM) channel pore gave rise to the voltage sensor domain (VSD) in voltage-gated potassium (Kv) channels and the voltage sensor-like domain (VSLD) in transient receptor potential (TRP) channels. Kv channels possess conserved positively charged residues in S4 (black plus signs), whereas TRP channels possess a TRP helix domain extending from the C-terminus of S6. Representative structures of the transmembrane domains of a 2TM channel (KcsA, PDB ID: 3eff, Uysal et al., 2009), a Kv channel (human Kv7.1, PDB ID: 6uzz, Sun and MacKinnon, 2020), and a TRP channel (rat TRPV1, PDB ID: 3j5p, Cao et al., 2013) are shown on the right, with one subunit of each tetramer displayed in color.

Six-transmembrane (6TM) ion channels are a major class of receptor proteins responsible for transducing signals into the output of ionic conductance across membranes. In ion channel gating, sensor activation opens or closes the pore, which is the effector domain of channel proteins (Fig. 1B). Structural homology suggests that 6TM channels emerged from the fusion of the four transmembrane helices (S1-S4) of the voltage sensor domain (VSD) to the pore of an ancestral two-transmembrane channel (Jegla et al., 2009) (Fig. 1C). Modern 6TM channels comprise several functionally and structurally distinct superfamilies (Yu and Catterall, 2004). Here, we compare two of these superfamilies, voltage-gated potassium (Kv) channels and transient receptor potential (TRP) channels, to look for common principles in the arrangement of their sensors and effectors. Kv channels primarily activate by sensing changes in membrane potential via a conserved set of positively charged residues, known as gating charges, in S4 of the VSD (Cui, 2016). In contrast, TRP channels lack conserved gating charges in the S1-S4 voltage sensor-like domain (VSLD) and acquired a C-terminal TRP helix domain extending from S6 (Venkatachalam and Montell, 2007). Members of the TRP superfamily have diverse sensory mechanisms and respond to stimuli of multiple modalities, including temperature, voltage, pH, mechanical force, and a wide array of ligands. Thus, Kv and TRP channels likely evolved divergent sensory capacities while maintaining a conserved effector domain.

To investigate the functional architecture underlying sensor evolution in 6TM channels, we analyzed the patterns of coevolution present at the superfamily, family, and subfamily levels in Kv and TRP channels using statistical coupling analysis (SCA). In brief, SCA computes the second-order correlations between pairs of residue positions in a multiple sequence alignment (MSA) to identify networks of mutually coevolving positions, known as sectors, within a group of homologous proteins (Halabi et al., 2009; Rivoire et al., 2016). In theory, sectors denote regions of the protein that coevolve due to shared functional constraints. Previous applications of SCA have validated this method for elucidating the structural units that mediate discrete functions in a variety of protein superfamilies, including serine proteases (Halabi et al., 2009), G-proteins (Rivoire et al., 2016), and protein kinases (Creixell et al., 2018).

In this study, we apply SCA to a large dataset of TRP and Kv channels to reveal the functional decomposition of 6TM channels into a ubiquitous “core” sector present at the superfamily level along with family-specific “accessory” sectors. By mapping existing structural and functional data of various channels in relation to the SCA results, we implicate the core sector in effector functions of the channel, such as pore gating and sensor-pore coupling. In contrast, we show that sensory structures, including ligand binding sites, voltage sensors, and modulators of temperature gating, correspond to accessory sectors and organize around the core sector according to three general principles that are conserved across TRP and Kv channels. These principles may reflect the functional constraints of sensor evolution around a conserved effector domain, providing an empirical paradigm of receptor protein architecture that may inform future approaches to protein engineering and rational drug design.

## Results

### Statistical coupling analysis of the TRP channel superfamily

To identify the sector decomposition of the TRP channel superfamily, SCA was performed on a manually curated multiple sequence alignment of the 7 major metazoan TRP families: TRPC, TRPM, TRPN, TRPA, TRPV, TRPP, and TRPML. The final alignment of the conserved transmembrane and TRP helix (S1-TRP) domains consisted of 1276 sequences comprehensively sampled from 56 species with phylogenetic representation from cnidarians to mammals (Table S1, Fig. 2A). Alignment processing identified 617 effective sequences (M’) and 242 positions with sufficient occupancy (≥80%) for statistical evaluation. Spectral decomposition and independent component analysis of the SCA covariance matrix identified 6 independent components (ICs), which comprise a total of 65 residue positions that display significant amino acid covariance when compared to the analysis of randomized alignments. Sectors were then identified by visual inspection as the largest combination of ICs showing mutual coevolutionary correlations, which are represented as warmer colors in the off-diagonal intersections of the IC submatrix (Fig. 2B). The resulting sectors were then projected onto the structure of rat TRPV1 (PDB ID: 3j5p, Cao et al., 2013), which we use as the reference structure for all TRP channel analyses.

**Figure 2.**
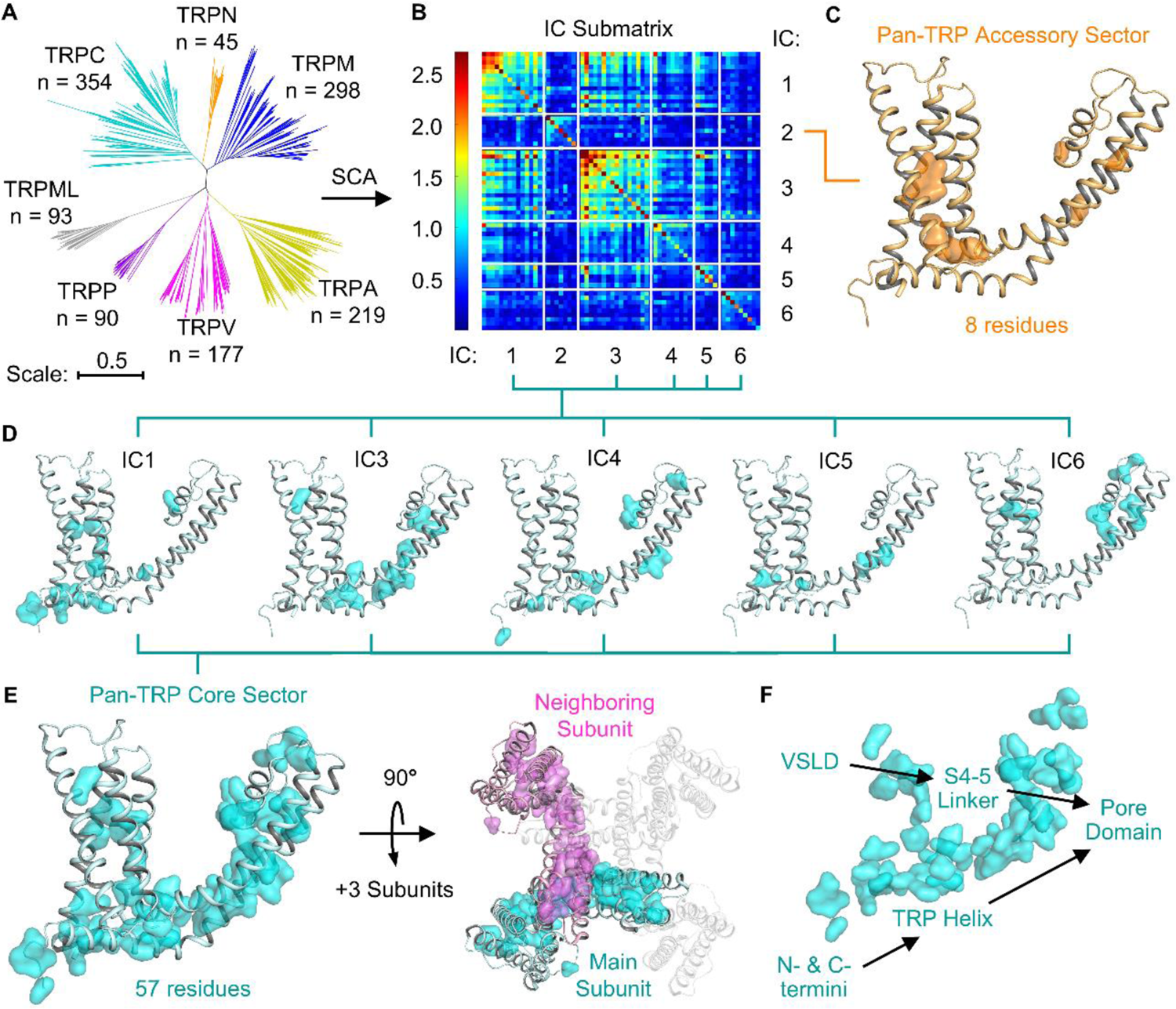
Statistical coupling analysis of the TRP channel superfamily. (A) Phylogenetic tree of the superfamily-wide TRP channel multiple sequence alignment. Clades representing the seven major TRP channel families (TRPC, TRPN, TRPM, TRPA, TRPV, TRPP, and TRPML) are shown in different colors. The number of sequence representatives from each family (n) is indicated. (B) Independent component (IC) submatrix depicting the amino acid covariation between residues within (on-diagonal boxes) and between (off-diagonal boxes) the six significant ICs identified by SCA. Warmer colors indicate a higher degree of covariance. (C) Structure of the pan-TRP accessory sector, corresponding to the residues in IC2 (orange surfaces), projected onto the reference structure of rat TRPV1 (PDB ID: 3j5p, Cao et al., 2013). (D) Structural projection of the ICs that show mutual inter-IC covariation (IC1 and IC3-6, cyan surfaces). (E) Projection of the pan-TRP core sector comprising IC1 and IC3-6 on a single subunit (left, cyan surfaces) and both the main (cyan) and neighboring (magenta) subunits of the channel tetramer (right). (F) The spatially contiguous organization of the core sector, which connects multiple channel domains to the pore. The cartoon backbone is omitted for clarity.

Upon projection, IC2 was found to consist of only 8 residues which exhibit minimal coevolution with all other ICs, indicating its identity as an independent protein sector. Due to its small size and coevolutionary independence, we define IC2 as the pan-TRP “accessory” sector. Positions defined by this sector were scattered throughout the channel subunit (Fig. 2C). Although the significance of small and distributed sectors remains unclear (Teşileanu et al., 2015), these residues were distributed at the points of closest contact between transmembrane helices (Fig. S1A), suggesting a possible role in the tuning of interhelical spacing and global channel conformation.

In contrast, the remaining five ICs (IC1 and IC3-6, Fig. 2D) were found to exhibit a high degree of mutual covariation, indicating their collective contribution to a single sector. We identify this sector as the pan-TRP “core” sector and primary sector of interest for three reasons. First, the sector is a contiguous network of 57 residues originating in the pore domain of the channel, occupying the interface between S5 and S6 as well as residues lining the ion conductance pathway (Fig. 2E). Hence, the core sector contains structures within the effector domain of the channel. Second, the sector connects the pore to other channel domains, including the VSLD and the N- and C-termini, through extensions along the S4-5 linker and TRP helix (Fig. 2F). Due to this structural connectivity, the core sector may mediate coupling between sensors and the pore domain, as indicated by our subsequent functional analyses. Third, because the SCA was performed on a representative superfamily-wide dataset, the core sector constitutes the largest coevolving unit that is conserved across all TRP channels. As such, this structure may have evolved as a result of the few functional constraints that are conserved across the diverse TRP superfamily, including pore gating, sensor-pore coupling, and permeation structure. Hence, the pan-TRP core sector may represent a conserved effector mechanism shared by TRP channels with divergent sensory functions.

### Organization of ligand binding sites around the pan-TRP core sector

Diverse ligands modulate TRP channel activation. To understand the relationship between TRP channel ligand sensitivity and the SCA results, we mapped all structurally resolved ligand binding sites in the transmembrane domain of TRP channels and analyzed their distribution relative to the pan-TRP core sector. A total of 57 ligands in 36 cryo-EM structures of ligand-bound TRP channels were assessed using LigPlot2.2+ (Laskowski and Swindells, 2011) to catalogue all amino acid positions involved in ligand-channel interactions, which were then mapped to their homologous positions in rat TRPV1. The resulting 108 unique positions were grouped into 9 distinct (but not mutually exclusive) binding sites harboring one or more ligands across different TRP channels (Table S2). This process is illustrated for site 1, which was identified by structural alignment of ligand-bound TRPV1, TRPV2, TRPV5, and TRPP1 channels (Fig. 3A). This site, which contains the vanilloid binding pocket in TRPV channels, was found to occupy 17 positions in S3, S4, the S4-5 linker, and the TRP domain of one subunit as well as three positions in the pore of the neighboring subunit (Fig. 3B, Table S2).

**Figure 3.**
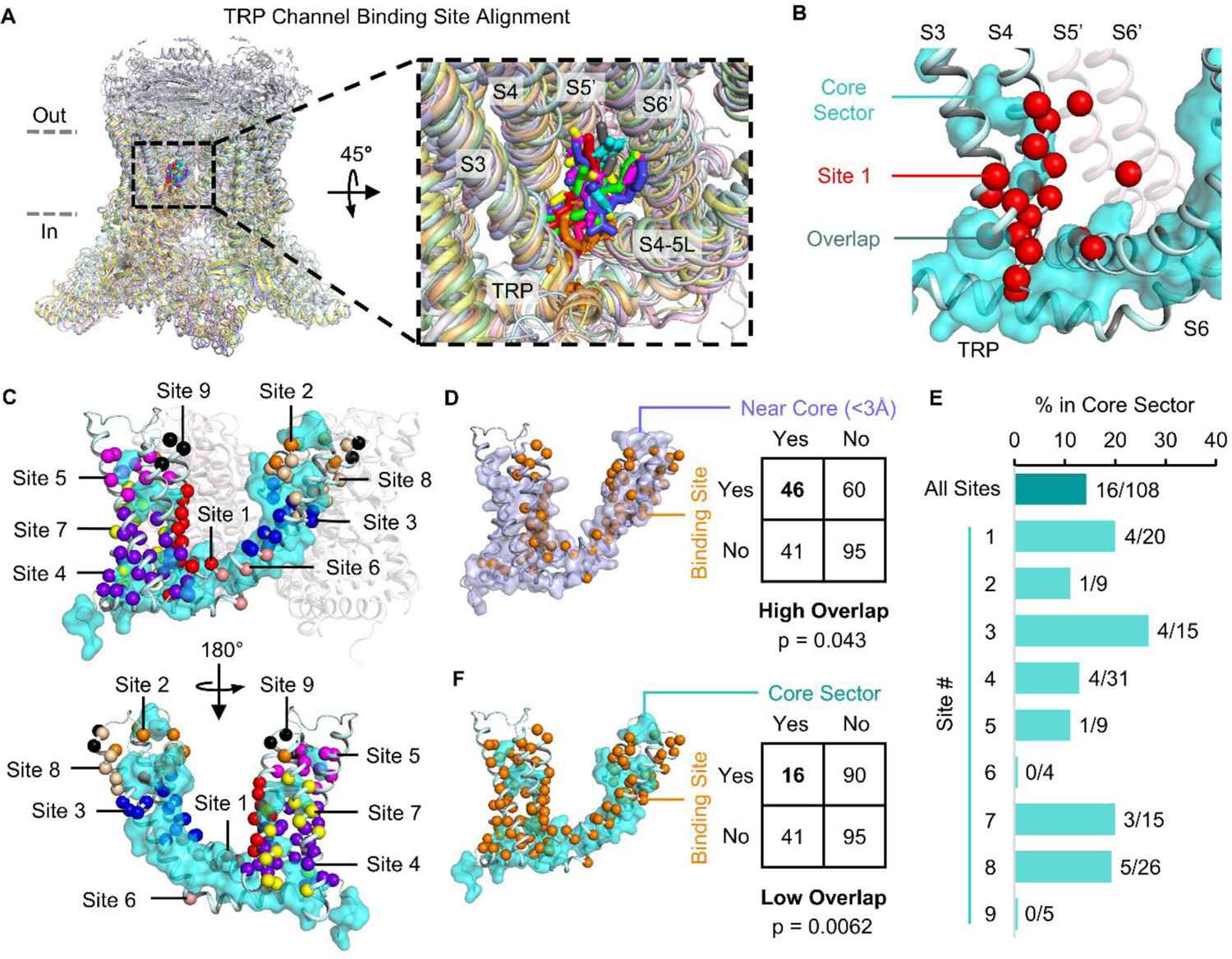
Organization of ligand binding sites around the pan-TRP core sector. (A) Structural alignment of TRP channels harboring ligands at site 1 (PDB IDs: 5IRX, 5IS0, 3J5Q, 6OO3, 6OO5, 6BWJ, 6B5V, 5IRZ, 6T9N, 5MKF) (See Table S2). Ligands are shown as sticks with different colors for each structure, while the cartoon channel structures are shown with corresponding light colors for clarity. (B) Residues forming site 1 (red spheres) projected with the pan-TRP core sector (cyan surface) onto the structure of rat TRPV1 (PDB ID: 3j5p, Cao et al., 2013). The neighboring subunit is shown in pink. (C) Overview of the 9 ligand binding sites and the pan-TRP core sector. Sites 1-9 are shown in different colors, with positions belonging to multiple sites shown in only one color for clarity. (D) Structural projection (left) and two-way contingency table (right) showing the distribution of residues between binding sites (orange spheres) and positions within 3 Å of the core sector (light blue surfaces). Bolded values indicate a higher degree of overlap than expected by random chance. (E) Proportion of overlapping positions between binding sites and the core sector. The fraction of binding site residues overlapping with core sector residues is indicated to the right each bar. (F) Structural projection (left) and two-way contingency table (right) showing the distribution of residues between binding sites (orange spheres) and the pan-TRP core sector (cyan surfaces). Bolded values indicate a lower degree of overlap than expected by random chance. For (D) and (F), p-values are calculated using the two-tailed Fisher’s exact test.

Binding site positions were found to be widely distributed across the channel subunit, occupying regions in the pore, VSLD, and TRP helix surrounding the core sector (Fig. 3C). This distribution suggests that ligand interactions can modulate the pore from nearly any region of the channel but are not systematically organized relative to any previously characterized structures in TRP channels.

Despite their diverse localization, all ligand binding sites were observed to reside near the pan-TRP core sector. We found that 43.4% of the binding site positions (46 of 108) were within 3 Å of the core sector (Fig. 3D). The observed proportion is significantly higher (p < 0.05, Fisher’s exact test) than the proportion expected by random chance (the “expected” proportion), which equals the total fraction of positions within 3 Å of the core sector (36%, or 87 of 242 total positions). Hence, binding site positions tend to occupy regions within molecular interaction distance of the pan-TRP core sector, suggesting that direct contact with the core is a constraint on ligand sensor localization.

Although binding sites emerge adjacent to the pan-TRP core sector, the extent to which they overlap with the core sector is limited. Individually, each site shares fewer than 6 positions with the core sector, with most sites having between one and five residues overlapping with the sector (Fig. 3E).

Across all sites, only 16 of 108 binding site positions (15.1%) were located within the pan-TRP core sector, which is significantly lower (p < 0.01, Fig. 3F) than the expected proportion (23.6%) given by the size of the sector (57 of 242 total positions). Thus, ligand binding sites predominantly occupy non-core positions in TRP channels.

Together, these results reveal that ligand sensors in TRP channels systematically organize relative to the core sector rather than conventional domains defined by structural homology. This sensor-core organization is governed by three general principles. First, sensors may emerge in any region surrounding the core. Hence, the core sector may mediate coupling between multiple peripheral domains and the pore. Second, sensors tend to reside adjacent to the core. Thus, interaction between sensors and the core sector may be required for channel modulation. Third, sensors tend to not be a part of the core. This indicates that core sector positions mediate non-sensor functions under strong evolutionary constraint, corroborating its role as an effector domain. As such, these three principles of sensor-core organization not only describe the superfamily-wide distribution of ligand sensors in TRP channels but also implicate the core sector in conserved coupling and effector functions.

### Sensor-core organization in the thermosensitive TRPV1-4 family

Next, we asked whether the core sector identified by the pan-TRP SCA is conserved in individual TRP families and subfamilies as well. In other words, is the core sector present and under strong coevolutionary constraint across phylogenetic scales? To explore this, SCA was performed on the vertebrate TRPV1-4 channels (2029 sequences, M’=130), which are the four subfamilies of the TRPV family that exhibit temperature-dependent activation (Benham et al., 2003) (Fig. 4A). The homology among these subfamilies allowed us to simultaneously analyze the transmembrane and cytosolic domains by SCA, which yielded a total of 21 ICs (Fig. S2A). In the S1-TRP domain, SCA identified two structurally contiguous ICs, Core IC1 and Core IC2 (Fig. 4B), which exhibit mutual coevolutionary coupling (Fig. S2A, left). Together, these two ICs show significant positional homology with the pan-TRP core sector, containing 40 of the 57 pan-TRP core sector positions (70.2%, p < 10^-8^). Hence, we defined the union of these two ICs as the TRPV1-4 core sector. In the N-terminus, four other ICs were found to form distinct contiguous networks (Fig. 4B). These include one IC located in the pre-S1 membrane proximal domain (MPD IC) and three ICs in the distal ankyrin repeat domain (ARD IC1-3), while the remaining ICs contained small groups of non-contiguous residues with unclear significance (Fig. S2A, right). Notably, the contiguous ICs in TRPV1-4 were not distinguishable by first-order conservation (Fig. S2A, bottom), demonstrating that differences in coevolution drive the observed sector decomposition. As such, these results confirm that the core sector is a coevolutionary feature of TRPV1-4 channels and reveal the presence of several accessory sectors in the N-terminal domain.

**Figure 4.**
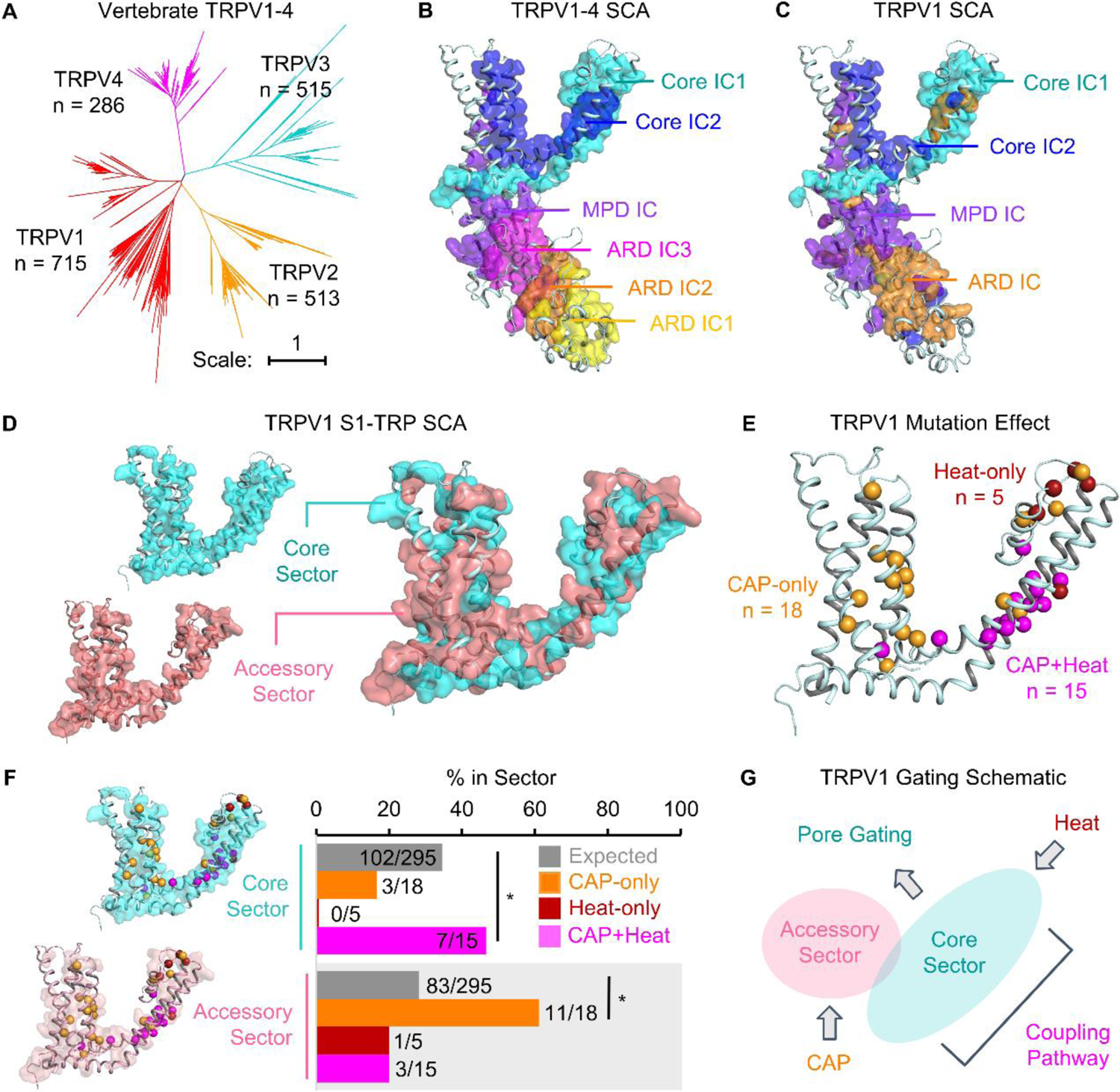
Sensor core organization in the thermosensitive TRPV1-4 channel family. (A) Phylogenetic tree of thermosensitive TRPV channels (TRPV1-4). Clades representing the TRPV1-4 subfamilies are shown in different colors. The number of sequences in each subfamily (n) is indicated. (B) Structurally contiguous ICs identified in the SCA of the TRPV1-4 family projected onto the structure of rat TRPV1 (PDB ID: 3j5p, Cao et al., 2013). (C) Structurally contiguous ICs identified in the SCA of the TRPV1 subfamily. (D) The TRPV1 core sector (cyan surfaces) and accessory sector (pink surfaces) identified in the SCA of the transmembrane and TRP (S1-TRP) domain of TRPV1 channels projected separately (left) and together (right). (E) Positions in TRPV1 that perturb channel responses to capsaicin only (CAP-only, orange spheres), heat only (Heat-only, red spheres), or both capsaicin and heat (CAP+Heat, magenta spheres) when mutagenized. (F) Structural projection (left) and percent overlap (right) of CAP-only, Heat-only, and CAP+Heat residues with the TRPV1 core and accessory sectors. The fraction of overlapping residues is given to the right of each bar. Expected values represent the percent overlap with the sector expected by random chance, which corresponds to the percent of the protein occupied by the sector (*p < 0.05, Fisher’s exact test). (G) Schematic of the relationship of capsaicin and heat stimulation to the core and accessory sectors in TRPV1 activation.

To probe smaller phylogenetic scales, SCA was performed on the TRPV1 channels as a subset of the TRPV1-4 dataset (621 sequences, M’=61, Fig. S2B). The resulting IC decomposition recapitulated that observed in TRPV1-4, identifying two Core ICs in addition to two N-terminal ICs representing combinations of the MPD and ARD ICs (Fig. 4C). To ensure that the observed similarities between the TRPV1-4 and TRPV1 SCA results were not driven by the TRPV1 subfamily alone, SCA was also performed on a dataset containing TRPV2-4 and mapped to the rat TRPV1 reference structure, which reproduced the results of the TRPV1-4 SCA (Fig. S2C). Hence, the TRPV core sector is conserved in but not exclusive to TRPV1 channels. Furthermore, when SCA was performed on the S1-TRP domain of TRPV1 (Fig. S2D), we identified an additional TRPV1 accessory sector (Fig. 4D) that was not present in a similar analysis of the S1-TRP domain in TRPV1-4. These results reveal that the core sector is a distinct coevolving motif at the superfamily, family, and subfamily levels in TRP channels. In contrast, accessory sectors were observed to be unique to certain phylogenetic subsets and may therefore represent derived units of channel evolution.

Next, we assessed the functional significance of the core and accessory sectors in TRPV1 channels by comparing the SCA results with a review of experimentally characterized mutations implicated in TRPV1 responses to capsaicin (CAP) or heat stimulation (Winter et al., 2013). Mutations in the S1-TRP domain were classified based on whether they perturb sensitivity to either stimulus discretely (CAP-only or Heat-only) or in concert (CAP+Heat) (Fig. 4E). When superimposed with the TRPV1 SCA results, CAP+Heat residues were found to be significantly enriched in the core sector (11 of 15 CAP+Heat positions, p < 0.01; Fig. 4F). Because mutations in the core disrupt activation by both stimuli, the core sector may contribute to effector functions required for downstream channel activation. These may include the coupling of sensor input to the pore domain as well as pore opening. In contrast, CAP-only residues were found to be significantly enriched in the TRPV1 accessory sector, which contained 11 of the 18 CAP-only positions (Fig. 4F, p < 0.01). The TRPV1-specific accessory sector not only harbored CAP-only residues in the vanilloid pocket (binding site 1) but also structurally distant CAP-only mutations in the extracellular pore domain and VSLD. Hence, the TRPV1 accessory sector may mediate vanilloid sensitivity and explain the disruption of capsaicin responses by mutations outside of the vanilloid binding site.

A similar analysis of chimera experiments in TRPV1-4 supports the role of accessory sectors in sensory functions. Residues required to transfer RTX sensitivity from TRPV1 to TRPV2-3 (Yang et al., 2016; Zhang et al., 2016, 2019) (Fig. S3A) were found to be significantly enriched in positions from the TRPV1 accessory sector (7 of 11 exchanged positions, p < 0.05), corroborating the contribution of the TRPV1 accessory sector to vanilloid sensitivity. Likewise, the near complete and exclusive exchange of the TRPV1-4 MPD IC (28 of 32 MPD IC residues, p < 0.001) was involved in transferring the temperature activation threshold and enthalpy of TRPV1 to TRPV2-4 (Yao et al., 2011) (Fig. S3B), suggesting that temperature activation is modulated by the MPD IC.

Together, these results indicate that the core sector is a conserved structure that mediates effector functions shared by all TRP channels, while accessory sectors correspond to portable, channel-specific sensory modules. Like the ligand sensors mapped in the pan-TRP dataset, the TRPV1 accessory sector and MPD IC occupy distinct regions of the channel. Yet, both remain proximal to the core sector, with 69% of TRPV1 accessory sector positions within 3 Å of the core sector and the MPD IC interfacing with core sector positions in the TRP helix. Importantly, both accessory sectors coevolve independently from the core sector, which is consistent with their role in non-effector functions. Thus, the properties of core and accessory sectors in TRPV1-4 reflect the three principles of sensor-core organization, suggesting that the core sector is an effector unit around which channel-specific sensory or modulatory sectors emerge.

### Conserved sensor-core organization in the Kv and TRP superfamilies

In contrast to the diverse sensory capacities of TRP channels, Kv channels are primarily activated by changes in membrane potential, which are detected by highly conserved residues in the VSD. Kv channels also exhibit selective conductance for potassium ions, which is mediated by the selectivity filter located C-terminal to the pore helix between S5 and S6. Because Kv and TRP channels likely evolved from the same ancestral channel (Fig. 1C), we examined if Kv channels share a similar sensor-core organization with TRP channels.

Like TRP channels, most members of the Kv superfamily assemble as tetramers in a “domain-swapped” conformation, in which the pore of one subunit is close to the VSD of an adjacent subunit (Fig. 1C, right). Hence, to facilitate comparisons between these two superfamilies, we applied SCA to the transmembrane region of domain-swapped Kv channels (Kv1-9) curated from a sequence alignment (1303 sequences, M’=180, Fig. 5A) of voltage-gated ion channels (PF00520) in the Pfam database (Finn et al., 2016). SCA identified 5 significant ICs which segregated into 2 sectors according to their mutual inter-IC correlations (Fig. S4A). When projected onto the reference structure of human Kv7.1 (KCNQ1), the first sector, which we refer to as the pan-Kv core sector, defined 53 residues that form a contiguous network of residues containing the S5-S6 interface, the S4-5 linker, and positions surrounding the center of the VSD (Fig. 5B, left). When compared by sequence alignment, the pan-Kv core sector was found to share a significant proportion of positions with the pan-TRP core sector (36%, 19 of 53 pan-Kv core residues, p < 0.01). Residues shared by the pan-Kv and pan-TRP core sectors mapped to the intracellular pore domain as well as the C-terminal S4-5 linker (Fig. 5D). In contrast, core sector positions that were unique to the pan-TRP or pan-Kv core sectors localized to the extracellular pore domain and VSD/VSLD (Fig. S4B-C).

**Figure 5.**
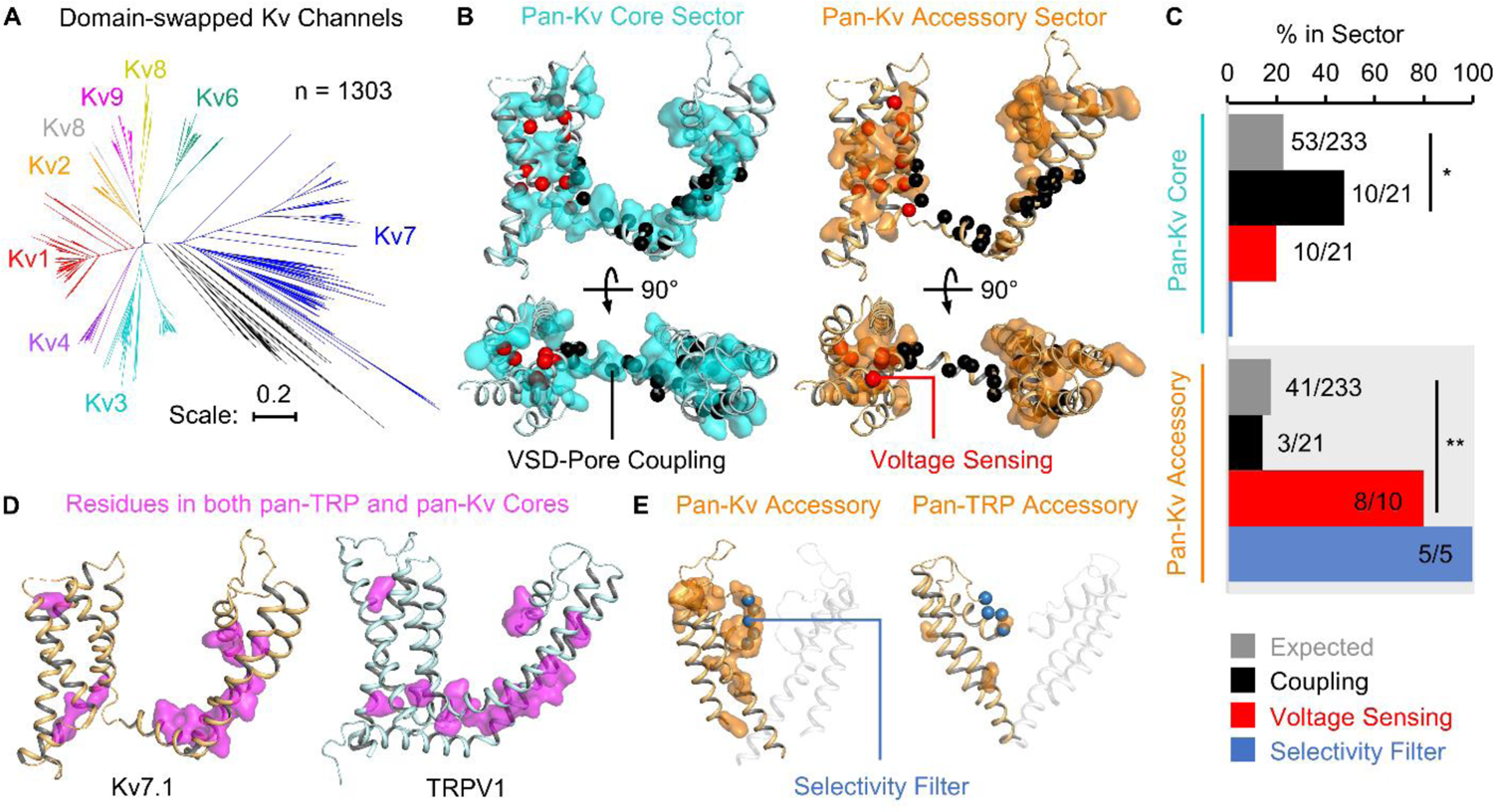
Conserved sensor-core organization in Kv and TRP channels. (A) Phylogenetic tree of domain-swapped Kv channels (Kv1-9). Clades representing each subfamily are shown in different colors. Black branches correspond to Kv channels with uncertain classification. The total number of sequences (n) is indicated. (B) Structure of the pan-Kv core sector (left, cyan surface) and pan-Kv accessory sector (right, orange surface) projected onto the structure of human Kv7.1 (PDB ID: 6uzz, Sun and MacKinnon, 2020). Residues implicated in VSD-pore coupling in Kv7.1 (black spheres) and voltage sensing (red spheres) are indicated. (C) Percent overlap of pan-Kv core and accessory sectors with residues involved in VSD-pore coupling, voltage sensing, and the selectivity filter in Kv7.1. Expected values represent the percent overlap with the sector expected by random chance, which corresponds to the percent of the protein occupied by the sector (*p < 0.05, **p < 0.01, Fisher’s exact test). (D) Projection of the 19 positionally homologous residues shared by the pan-Kv and pan-TRP core sectors (magenta surfaces) onto the structure of human Kv7.1 (left) and rat TRPV1 (right) (PDB ID: 3j5p, Cao et al., 2013). (E) Structural comparison of selectivity filter residues (blue spheres) in relation to the pan-Kv accessory sector (left) and pan-TRP accessory sector (right).

Previous studies in domain-swapped Kv channels have shown that the region shared between the pan-Kv and pan-TRP core sector is important for coupling between the VSD and the pore (Hou et al., 2020; Long et al., 2005; Lu et al., 2001, 2002) as well as pore opening (del Camino et al., 2000; Liu et al., 1997; Long et al., 2007). To examine whether the pan-Kv core sector is implicated in VSD-pore coupling, we compared the core sector with functionally significant residues in Kv7.1 channels. Kv7.1 is important for the function of cardiac and epithelial tissue and has been the subject of extensive structure-function studies (Cui, 2016). In prior work, we characterized 21 positions in Kv7.1 which mediate coupling between VSD activation and pore opening (Hou et al., 2020; Wu et al., 2010). Structural projection (Fig. 5B, left) showed that these coupling residues are significantly enriched in the pan-Kv core sector (10 of the 21 positions, 47.6%, p < 0.05) (Fig. 4C). Thus, like the pan-TRP core sector, the pan-Kv core sector harbors residues implicated in coupling sensory input to the channel pore, suggesting that the core sector is functionally similar in both superfamilies.

The second sector identified by SCA, which we define as the pan-Kv accessory sector, was found to comprise 41 residues which form two spatially distinct clusters adjacent to the pan-Kv core sector, one located in the center of the VSD and the other in the pore helix and the selectivity filter (Fig. 5B, right). In the VSD, multiple residues contribute to sensing membrane potential. These include six positively charged residues in S4 (R228, R231, Q234, R237, H240, and R243) (Bezanilla, 2000); positions in the charge transfer center that facilitate the movement of the charged residues in S4 across the membrane (F167, E170, and D202N) (Lacroix et al., 2014; Tao et al., 2010); and E160 in S2, which interacts with the charged residues in S4 during VSD activation (Wu et al., 2010). The pan-Kv accessory sector was found to contain 8 of these 10 residues (Fig. 5B-C), indicating that positions involved in voltage sensing are significantly enriched in this sector (p < 0.01). Hence, this sector may represent the coevolutionary unit mediating voltage sensitivity within the Kv superfamily.

Similar to the sensors observed in the TRP superfamily, the pan-Kv accessory sector emerges in close proximity to the pan-Kv core sector but remains functionally and coevolutionarily distinct from it. Within the VSD, the inwards-facing residues of the pan-Kv accessory sector interface extensively with the surrounding core sector positions, thereby enabling their direct interaction. Moreover, the lack of covariation between the two sectors (Fig. S4A, right) is consistent with their disparate functions, with the core sector contributing to coupling as well as pore opening and the accessory sector to voltage sensing. Thus, the voltage sensor in Kv channels organizes relative to the pan-Kv core sector according to the same principles observed in the TRP superfamily.

In addition to comprising voltage-sensing residues in the VSD, the pan-Kv accessory sector also occupies a cluster of positions in the selectivity filter of the pore domain. This region of the accessory sector was found to contain all 5 residues of the conserved T(V/I)GYG motif (T312-G316 in human Kv7.1) that defines the canonical potassium selectivity filter, indicating that the pan-Kv accessory sector mediates the ionic specificity of channel conductance (Fig. 5C, E).

The fact that both voltage sensitivity and potassium selectivity are mediated by a single sector may suggest that these two functions evolve under a common coevolutionary constraint within the Kv superfamily. However, because these positions are highly conserved in Kv channels, a broader dataset may be required to examine coevolution between the voltage sensor and selectivity filter in greater detail.

Like the voltage sensor, the selectivity filter of the pan-Kv accessory sector extends from the periphery of the core sector but demonstrates limited coevolutionary coupling with it. The principles of sensor organization relative to the core sector may therefore apply to other determinants of specificity in ion channels, including regions like the selectivity filter, which modify the output of the channel yet are not conserved across all channel types. In comparison, the pan-TRP accessory sector contains only one position in the selectivity filter corresponding to a highly conserved glycine (G643 in rat TRPV1) (Fig. 5E), which forms the narrowest region of the pore in TRPV1 (Cao et al., 2013). Thus, the pan-TRP accessory sector is greatly diminished in size and functional specificity relative to the pan-Kv accessory sector, which is consistent with the diversity of sensors and lack of cation selectivity that characterizes the TRP superfamily.

### Sensor-core organization governs the location of pathogenic substitutions, exchangeable domains, and artificial ligand interactions

Coevolving networks of core and accessary sectors are central to sensor-effector organization, but can these sectors explain the segregation of other functionally important positions in channel proteins? To explore this, we compared the results of SCA with pathogenic substitutions, chimera experiments, and the binding sites of an artificial ligand in TRP and Kv channels.

First, we studied the distribution of clinically characterized missense variant sites in human TRP and Kv channels. In the S1-TRP domain of human TRPV1-4, a total of 20 pathogenic and 12 benign variants were reported in the ClinVar database (Landrum et al., 2018) (Fig. 6A, left). The core sector formed by Core IC1 and Core IC2 was found to be significantly enriched in pathogenic variants, containing 17 of the 20 sites (85%, p < 0.001) (Fig. 6A, right). Likewise, at the superfamily level, the pan-TRP core sector was found to be significantly enriched in pathogenic variants from all human TRP channels (p < 0.01), while pathogenic variants in human Kv1-9 were significantly overrepresented in both the pan-Kv core (p < 0.05) and accessory (p < 0.01) sectors (Fig. S5A-B). In contrast, benign variant sites showed no significant association with any sectors (p > 0.05). Hence, core and accessory sectors are enriched in functionally important residues that are perturbed by disease-associated mutations across diverse channel families.

**Figure 6.**
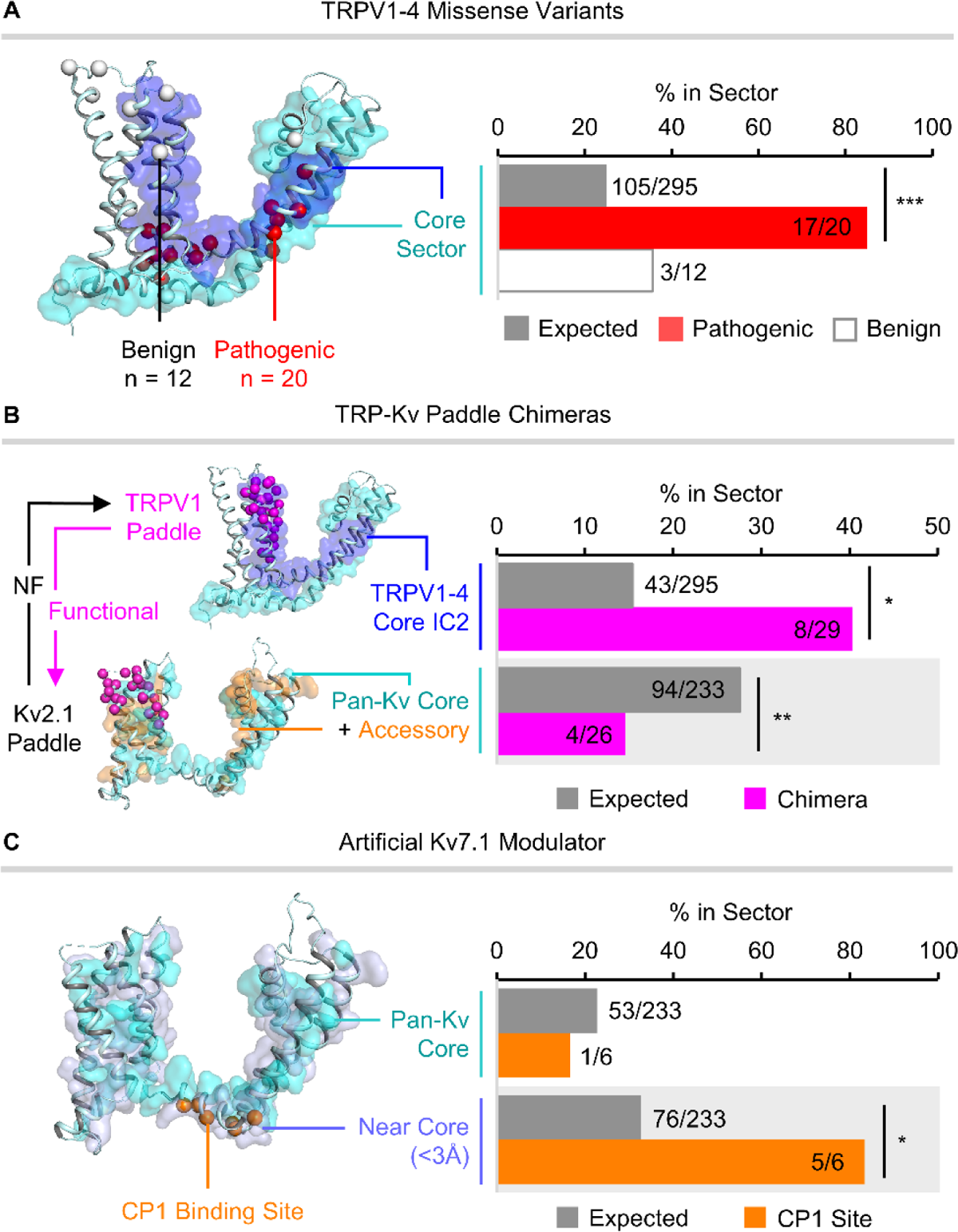
Sensor-core organization governs the location of pathogenic substitutions, exchangeable domains, and artificial ligand interactions. (A) Structural comparison (left) and percent overlap (right) of the TRPV1-4 core sector (Core IC1, cyan surface; Core IC2, blue surfaces) with the 20 pathogenic (red spheres) and 12 benign missense variants sites identified in human TRPV1-4 channels. (B) As in (A), comparison of the TRPV1-4 Core IC2 (blue surface) and the pan-Kv core (cyan surfaces) and accessory (orange surfaces) sectors with a region of the S3-S4b paddle motif (magenta spheres) that produced functional chimeras when transferred from TRPV1 to Kv2.1 but not in the reciprocal exchange from Kv2.1 to TRPV1. (C) As in (A), comparison of the pan-Kv core sector (cyan surfaces) and residues within 3 Å of the pan-Kv core sector (light blue surfaces) with CP1 binding site residues (orange spheres; R249, L250, S253, Q260, K354, V355, and K358) in human Kv7.1. K358 (not shown) lacked sufficient conservation to be analyzed by SCA and was therefore excluded from the analysis. Rat TRPV1 (PDB ID: 3j5p, Cao et al., 2013) and human Kv7.1 (PDB ID: 6uzz, Sun and MacKinnon, 2020) are used as the reference structures for all TRP and Kv channel projections, respectively. Expected values represent the percent overlap with the sector expected by random chance, which corresponds to the percent of the protein occupied by the sector (*p < 0.05, **p < 0.01, ***p < 0.001, Fisher’s exact test).

Next, to determine the relevance of sectors to structure-function studies of ion channels, we examined a chimera study which systematically exchanged transmembrane region between Kv2.1 and two TRP channels, TRPV1 and TRPM8. Here, functional (conductive) hybrid channels were only obtained when the S3b-S4 paddle motif of Kv2.1 was replaced by the homologous paddle motif of TRPV1 or TRPM8 (Kalia and Swartz, 2013). Importantly, the reciprocal paddle chimeras in both TRP backgrounds were non-functional. To understand the unidirectional success of this chimera experiment, we compared the largest reciprocally tested S3b-S4 paddle motif that produced a functional chimera in the Kv2.1 but not TRPV1 background (2V1Kv) with the TRPV1 and pan-Kv SCA results (Fig.6B). The TRPV1 paddle (V525-L553) was found to be significantly enriched in Core IC2 positions (p < 0.05), whereas the Kv paddle (V211-L236 in Kv7.1) was significantly depleted in pan-Kv core and accessory sector positions (p < 0.01). Hence, the paddle motif may be asymmetrically intolerant of the substitution because it harbors a large component of the core sector in TRPV1 but not Kv2.1. Other chimera experiments also suggest that an intact core sector is important for channel function. In the RTX sensor and MPD chimera experiments described previously (Fig. S3A-B), the exchanged regions contained fewer than 3 core sector positions each, indicating that the core sector remained unperturbed in both functional hybrids. Moreover, two segments of the S4-5 linker and S6 which are jointly required to establish coupling between the Shaker (Kv1.2) VSD and two-transmembrane KcsA channel were found to map to the pan-Kv core sector (Fig. S5C), corroborating the role of the core sector in VSD-pore coupling. As such, successful chimera experiments appear to obey the constraints of sensor-core architecture.

Lastly, we analyzed the binding site of CP1, a synthetic compound resembling the head group of phosphatidylinositol-4,5-bisphosphate (PIP2) that was found to enhance VSD-pore coupling in Kv7 channels (Liu et al., 2020). In human Kv7.1, the binding site of CP1 comprises seven residues (R249, L250, S253, Q260, K354, V355, and K358) which line the S4-5 linker and S6 (Fig. 6C, left). One position, K358, was found to be within 3.3 Å of the pan-Kv core sector but lacked sufficient homology across Kv channels to be analyzed by SCA. The remaining 6 residues were found to be significantly enriched among non-core positions within 3 Å of the pan-Kv core sector (p < 0.05, Fig. 6C, right), with 5 positions (83.3%) located directly adjacent to the core sector. Similarly, the purported PIP2 binding site targeted by CP1 also localizes to non-core positions near the pan-Kv core sector (Fig. S5D). Hence, binding sites in Kv channels, like those in the TRP superfamily, localize according to the same principles of sensor-core organization. Together, these results demonstrate that the constraints of sensor-core architecture reflect the structure-function relationships observed in 6TM channels across a variety of datasets.

## Discussion

As revealed through SCA of the TRP and Kv superfamilies, 6TM channels share a general coevolutionary organization that underlies their capacity to transduce diverse stimuli to the output of ionic conductance. Central to this architecture is the core sector, which emerges as the coevolutionary network uniting channels of divergent sensory capacities. In both TRP and Kv channels, the core sector takes the form of a sparse yet contiguous network of positions originating and extending from the channel pore that exhibit consistent patterns of coevolution at the subfamily, family, and superfamily levels. Functional studies in individual channels, including TRPV1 and Kv7.1, implicate the core in downstream effector channel functions, such as pore gating and sensor-pore coupling (Fig. S6A). Furthermore, the analysis of disease-associated substitutions and channel chimeras both indicate that an intact core sector is essential for baseline channel activity. Due to its conservation and functional significance, the core sector may represent the effector of domain-swapped 6TM channels that contains the basic scaffold and actuation mechanisms necessary for gated ionic conductance.

Around the core sector, sensors responsible for detecting stimuli of various modalities are arranged according to three fundamental principles. First, sensors may emerge in nearly any region of the channel with access to the core. From ligand binding sites in TRP channels to voltage sensing structures in the Kv VSD, residues mediating stimulus detection occur in structurally diverse areas, suggesting that the core sector mediates allosteric coupling between various sensory regions to the pore. Second, sensors must maintain contact with the core sector. Proximity to the core may be required for the physical transduction of sensor movement to the coupling network within the core sector, which then opens or closes the channel pore. Third, sensors must not disrupt the integrity of the core. As observed in the analysis of pathogenic substitutions and channel chimeras, residues within the core sector are functionally intolerant of substitutions, which may restrict the emergence of sensors to non-core regions. Together, these three principles of sensor-core organization capture the internal architecture shared by functionally and evolutionarily disparate channels, providing an empirical description of the higher-order structure present within a major class of receptor proteins.

These three principles of sensor-effector organization point to a plausible model of sector evolution within the established evolutionary history of 6TM channels (Fig. 1C). Due to its strong conservation, the core sector motif may represent a basic effector unit originating in the ancestral 2TM channel. The addition of the VSD in 6TM channels may have driven the development of the core sector to accommodate coupling to the S1-S4 domain, resulting in the extension of the core along the S4-5 linker observed in both TRP and Kv channels. In TRP channels, the acquisition of the TRP helix may have enabled a similar expansion of the core sector to couple diverse input from both the VSLD and intracellular domains. While the core sector has remained ubiquitously conserved, family-specific accessory sectors and sensors may emerge or fade in order to fulfill the functional demands of their specific cellular contexts. In Kv channels, selective pressures for voltage sensitivity and potassium selectivity likely gave rise to the pan-Kv accessory sector, which comprises regions in the VSD and selectivity filter. In contrast, the absence of such pressures may have diminished coevolution within the VSD and selectivity filter in TRP channels, facilitating the diversification of stimulus sensors in unconstrained regions adjacent to the core sector to produce the diverse repertoire of sensory capacities observed in the modern superfamily.

Consistent with the results of SCA in other protein families, the coevolutionary analysis of ion channels reveals a new paradigm of the structure-function relationships within receptor proteins. In contrast to conventional domains defined by structural homology, the core-accessory model of channel sectors identifies the internal networks of residues responsible for conserved or specific receptor functions (Fig. S6A) and provides generalizable principles describing their arrangement. Because the three principles of sensor-core organization likely result from an evolutionary pattern of sensor diversification around a conserved effector, the same principles may apply to other receptor proteins that emerged from a conserved output modality, including the G-protein coupled receptors, receptor kinases, and nuclear receptors. As such, the conserved principles of sensor-core organization in 6TM channels may provide new insights into the architecture of signal transduction in a wide array of receptor proteins and inform future approaches to protein engineering and rational drug design.

## Acknowledgements

We thank Dr. Lawrence B. Salkoff for a critical reading of the manuscript and helpful suggestions. This work was supported by National Institutes of Health Grants R01MH116981 (to J.C.) and R01GM132110 (to J.Z.).

## Author contributions

Conceptualization, J.C.; Methodology, A.D., P.W.K., P.H., T.J., M.A.Z., and J.Z.; Investigation, Data Curation and Visualization, A.D., P.W.K., and P.H.; Writing – Original Draft, A.D. and J.C.; Writing– Review & Edit, A.D., P.W.K., P.H., M.A.Z., J.Z., T.J., and J.C.

## Methods

### Multiple sequence alignment generation

To generate an alignment of the TRP channel superfamily, sequences from members of the seven major TRP channel subfamilies (TRPC, TRPM, TRPN, TRPA, TRPV, TRPP, and TRPML) were comprehensively collected from 56 metazoan species (Table S2) using protein-to-nucleotide BLAST searches in the NCBI RefSeq RNA database (O’Leary et al., 2016). The TRPY family was excluded because it is exclusive to yeasts, and the recently identified TRPS family, a sister clade of TRPM that is largely restricted to protostomes, was grouped within the TRPM family. Query sequences comprised human TRPC, TRPM, TRPA, TRPV, TRPP, and TRPML and Drosophila melanogaster TRPN1 (NOMPC). Homologs were classified by TRP subfamily using NCBI annotations, back-BLAST searches in mouse and zebrafish, and the presence of characteristic sequence motifs.

All sequence alignments were performed using ClustalOmega (Sievers and Higgins, 2014). To produce a superfamily-wide alignment, we first generated alignments of each TRP family, which were then truncated to the transmembrane domain and TRP helix domains using representative TRP channel structures as a guide. In the case of TRPP and TRPML, the lipase domain, corresponding to R251-G449 in TRPP1 and T121-A294 in TRPML1, was excluded due to their lack of conservation in other TRP channel families. Then, the truncated sequences were combined and realigned to generate the superfamily-wide alignment. This alignment was then manually edited to remove highly gapped sequences, sequences with unresolved residues designations, and excessive gaps within transmembrane segments. The final alignment consisted of 1276 sequences comprising 354 TRPC, 298 TRPM, 45 TRPN, 219 TRPA, 177 TRPV, 90 TRPP, and 93 TRPML orthologs.

For the analysis of temperature sensitive TRPV channels, all vertebrate TRPV1-4 sequences were collected from the NCBI Identical Protein Groups database (Benson et al., 2018) by keyword searches following the NCBI standard annotation of TRP channels (e.g. “transient receptor potential cation channel subfamily V member 1” for TRPV1). These sequences were then filtered to exclude sequence fragments and sequences with excessively long insertions, resulting in a total of 2029 sequences representing 715 TRPV1, 513 TRPV2, 515 TRPV3, and 286 TRPV4 channels. These sequences were then aligned to produce the full length TRPV1-4 alignment. To derive the alignments of TRPV1 and TRPV2-4, the associated sequences were extracted from the TRPV1-4 alignment without further realignment. Likewise, the full length TRPV1 alignment was truncated to positions corresponding to K425-A719 in rat TRPV1 for the analysis of the transmembrane and TRP domains of TRPV1.

The alignment of domain-swapped Kv channels was derived from the PF00520 alignment of 6TM ion channel transmembrane domains obtained from the Pfam database (Finn et al., 2016). Sequences of domain swapped Kv channels, corresponding to Kv1-9, were extracted by Pfam annotation, as described previously (Hou et al., 2020), and any highly gapped or poorly aligned sequences were removed. The final alignment consisted of 1303 Kv1-9 transmembrane domain sequences.

### Phylogeny construction

Phylogenetic trees of the pan-TRP, TRPV1-4, and pan-Kv alignments were constructed using the neighbor-joining method implemented in MEGA (Kumar et al., 2018). Branch lengths represent Poisson Correction distance.

### Statistical coupling analysis

SCA was performed using the pySCA software package (Rivoire et al., 2016). Detailed descriptions of the SCA calculations are reported in prior work (Halabi et al., 2009; Rivoire et al., 2016). In brief, SCA computes a conservation-weighted covariance matrix for all pairwise positions in a multiple sequence alignment that is processed to remove highly gapped positions and sequences. This covariance matrix is then subjected to eigenvalue decomposition to identify the top eigenmodes corresponding to the positions exhibiting the greatest covariance in the dataset. Independent component analysis transforms the top eigenmodes into independent components (ICs), which represent groups of mutually covarying positions which exhibit minimal inter-group covariance. The top-contributing residues in each IC are then selected from the cumulative density function according to a user-defined confidence threshold, which is set to 0.95 by default. The resulting positions in each IC are plotted in an IC submatrix, in which the intensities of the on-diagonal boxes represent the strength of intra-IC correlations, with those of the off-diagonal boxes representing the strength of inter-IC correlations. Sectors are defined as the largest linear combinations of ICs which exhibit mutual correlations within the IC submatrix by inspection. The resulting sector and IC positions are then mapped to positions in a reference sequence for structural projection.

For this analysis, rat TRPV1 is used as the reference structure for all TRP channel SCA results, while human Kv7.1 is used as the reference structure for the SCA of Kv channels. Alignment processing parameters, effective sequence numbers (used to calculate sequence weights), and the confidence thresholds used for selecting the top contributing residues in each IC are summarized for each SCA in Table S3.

### Ligand binding site determination

Ligand binding sites were obtained via analysis of 57 ligands in 36 cryo-EM structures of ligand-bound TRP channels. Ligands forming interactions with non-transmembrane regions of the channel were excluded. For each structure, binding site residues were determined using LigPlot2.2+ (Laskowski and Swindells, 2011) with default settings, which identifies all residue positions involved in hydrogen bonds or non-bonded contacts with the ligand. Residues forming hydrogen bonds were defined by a minimum hydrogen-acceptor distance of 2.7 Å and a maximum donor-acceptor distance of 3.3 5Å. Non-bonded contacts were defined as having a minimum contact distance of 2.9 Å and a maximum contact distance of 3.90 Å. Binding site residues were then grouped into binding sites according to the proximity of ligands in structural alignments of the transmembrane domain of different ligand-bound channels. Ligands that were visibly overlapping in the structural alignment were grouped into the same binding site, while non-overlapping ligands were grouped into separate sites. All binding site residues were then mapped to their homologous positions in the sequence of rat TRPV1 via multiple sequence alignment. Six binding site residues that occupied positions that were not analyzed in the SCA due to poor alignment quality were excluded in this step. The final TRP channel binding sites comprised all unique residues identified in this manner. In human Kv7.1, residues involved in the binding site of CP1 (R249, L250, S253, Q260, K354, V355, and K358) and PIP2 (F127, F130, G246, T247, R259, Q260, I263, T264, K354, and K358) were identified by molecular docking and mutagenesis in a previous study (Liu et al., 2020). Residues F127 and K358 lacked sufficient homology to be analyzed by SCA and were therefore excluded from sector comparisons.

### Classification of TRPV1 mutagenesis data

Functionally significant residues in TRPV1 identified by mutagenesis were obtained from a comprehensive review of experimental studies conducted before 2013 (Winter et al., 2013). Of the 112 total mutated sites, analysis was restricted to the 79 characterized residues within the transmembrane and TRP domains (K425-A719) of TRPV1. Of these, 38 residues which were implicated in channel activation by capsaicin or heat were identified according to the annotations included in the review, which denote positions in which mutations cause measurable changes in the activation of TRPV1 in response to either stimulus. This includes shifts in current responses induced by capsaicin or heat, the potentiation of capsaicin-induced currents by heat, and the potentiation of heat-induced currents by capsaicin. These residues were then further classified based on whether their mutation influenced both capsaicin and heat responses simultaneously (CAP+Heat), capsaicin responses alone (CAP-only), or heat responses alone (Heat-only). These three categories, which are mutually exclusive and comprehensive, were then used in comparisons with sector positions.

### First-order amino acid conservation

The first-order conservation of amino acids was calculated using the Kullback-Leibler relative entropy (Di), which quantifies the difference between the observed frequency of residues at a position to the background amino acid frequencies in a sequence dataset. To compare the conservation of residues in the TRPV1-4 ICs, the average and standard deviation of the Di was calculated for all positions in each IC.

### Clinical missense variant analysis

Positions in human TRP channels associated with missense variants of known clinical significance were obtained from the NCBI ClinVar database (Landrum et al., 2018). The residue positions associated with single gene missense variants annotated as either pathogenic or benign according to standard guidelines (Richards et al., 2015) were comprehensively catalogued for each member of the human TRP channel family through database searches of the corresponding genes. Surveyed channels included human TRPC1, TRPC3-7, TRPM1-8, TRPA1, TRPV1-6, TRPP1-3, and TRPML1-3, although no variants meeting the above criteria were reported for TRPV2, TRPC1, TRPC4-7, TRPML3, and TRPP3. Residue positions associated with each variant were then mapped to their corresponding positions in rat TRPV1, and any positions which were not within the transmembrane and TRP domains were excluded. The final dataset identified a total of 30 pathogenic and 20 benign missense variants across all TRPs, of which 20 pathogenic and 12 benign variants were present in the TRPV family. To identify clinical missense variants in the human Kv superfamily, the same process was applied to the human set of domain-swapped Kv channels comprising Kv1-9. Variant positions in each channel were then mapped to residues in Kv7.1, which resulted in a total of 116 pathogenic and 8 benign sites.

### Chimera analysis

All chimera positions were identified from prior studies and mapped to the corresponding positions in rat TRPV1 by sequence alignment. Positions involved in transferring RTX sensitivity from TRPV1 to TRPV2 and TRPV3 were obtained from three studies: The first identified four positions in mouse TRPV1, corresponding to S512, F543, T550, and E570 in TRPV1, which produced partial RTX-dependent activation when transplanted to homologous positions in TRPV2 (Yang et al., 2016). A subsequent study demonstrated that transferring S512, M547, T550, and E570 produced maximal RTX sensitivity in a rat TRPV2 chimera (Zhang et al., 2016). Hence, F543 was excluded from this set. A third study found that six mutations were sufficient to produce partial activation of TRPV3 by RTX, which comprised Y511 and E513 and the four previously identified residues (Zhang et al., 2019). However, robust activation required an additional five residues in the pore domain, L577, V596, L630, I661, and Y671. Thus, our analysis included all eleven of these positions (Y511, S512, E513, M547, T550, E570, L577, V596, L630, I661, and Y671).

For the analysis of temperature activation chimeras, the membrane proximal domain comprised the segment of TRPV1 (H358-F434) exchanged in the original study (Yao et al., 2011), which was found to confer the temperature-current relationship of the donor channel when transplanted in chimeras of TRPV1-4. Our analysis describes the transfer of the TRPV1 MPD and temperature activation properties to other channels, as this was most extensively tested donor-recipient permutation reported in the study.

The segments of the S4-5 linker and S6 required for coupling the Shaker VSD to the KcsA pore domain were defined according to the complementary sequence motifs identified in Shaker, which are “LGRTLKASMRELGLL” and “PVPVIVSNFNYFY” (Lu et al., 2002). Upon alignment, these sequences map to L385-L399 and P473-Y485 in Kv7.1, which were the segments included in the analysis.

The results of chimeras between Kv and TRP channels were obtained from a study involving reciprocal exchange of multiple domains between Kv2.1 and either TRPV1 or TRPM8 (Kalia and Swartz, 2013). Non-functional chimeras refer to channel hybrids that are non-conductive in all tested conditions, whereas functional chimeras exhibit some measurable conductance constitutively or in response to activating stimuli. Of the 50 chimera permutations tested, 10 hybrid channels were classified as functional. All 10 functional chimeras involved transfer of portions of the S3b-S4 paddle motif from TRPV1 or TRPM8 to Kv2.1. For comparison, we restrict our analysis to the largest region of the paddle motif that produced a functional chimera when transplanted from TRPV1 to Kv2.1 (2V1Kv) but not when reciprocally transferred from Kv2.1 to TRPV1 (1KvV1), which corresponds to V211-L236 in Kv7.1 and V525-L553 in TRPV1.

### Statistical tests of association

All statistical tests of association were performed using the two-tailed Fisher’s exact test. The appropriate two-by-two contingency tables were constructed to represent the distribution of all analyzed residue positions between the sector and experimental categories. Unless otherwise stated, the total positions represent all positions in the channel region in which the SCA was performed.

## Supplemental figures with titles and legends

**Figure S1.**
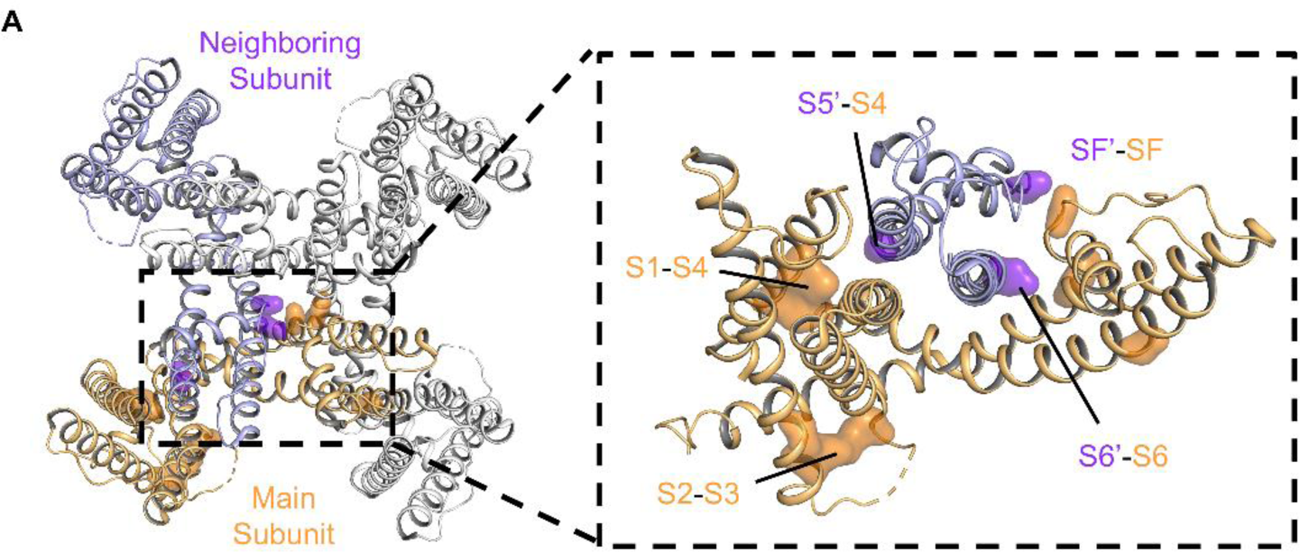
Distribution of accessory sector residues at interhelical contact points. (A) Projection of the pan-TRP accessory sector (surfaces) onto the structure of rat TRPV1 (PDB ID: 3j5p, Cao et al., 2013). The eight residues defining the pan-TRP accessory sector localize to the points of closest contact between helices (inset) within the main subunit (orange) and at interfaces with the neighboring subunit (purple).

**Figure S2.**
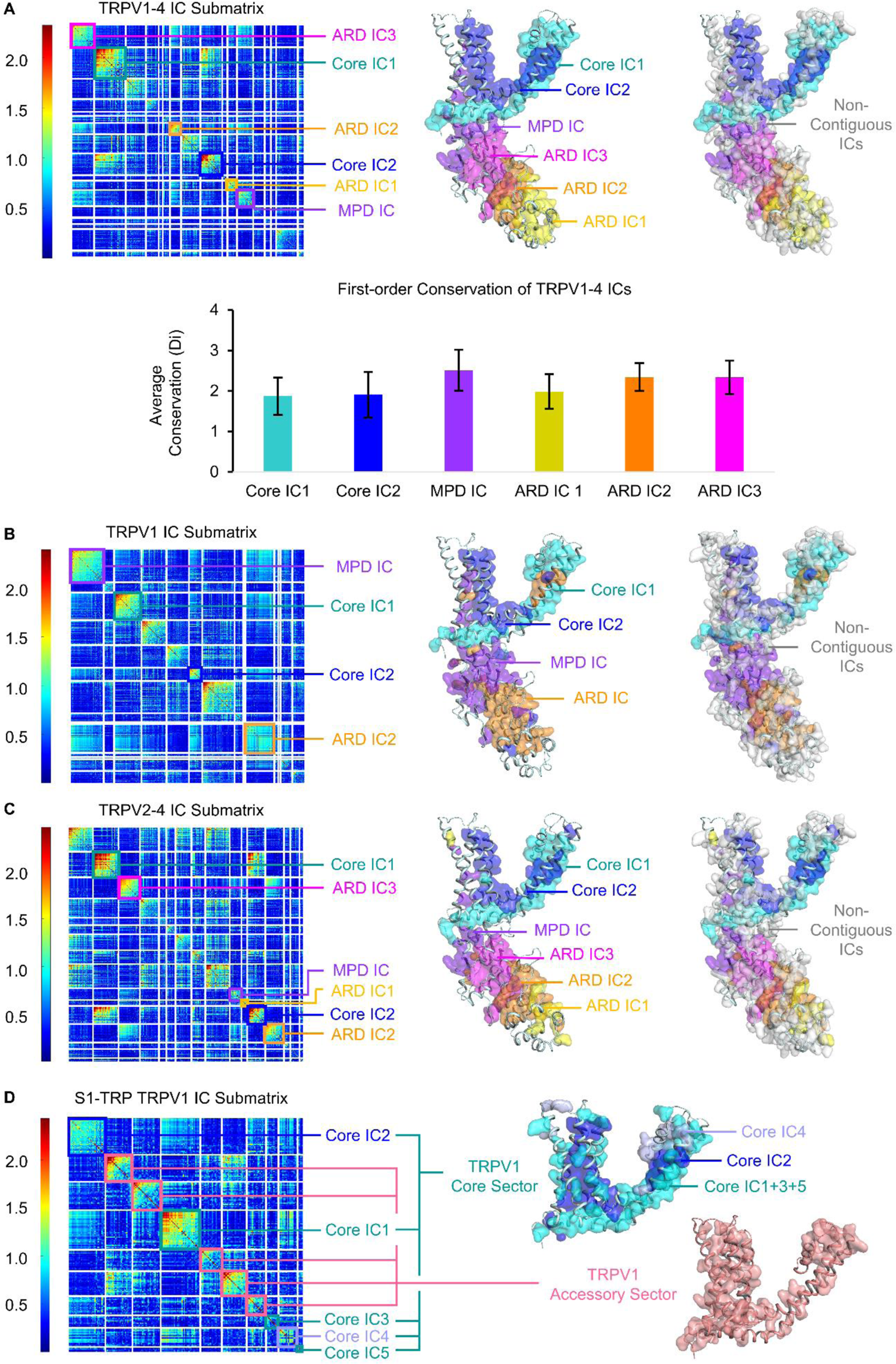
Sector decomposition of the TRPV1-4 family. (A-D) Independent component (IC) submatrices (left) and structural projections (rat TRPV1, PDB ID: 3j5p, Cao et al., 2013) of the contiguous (middle) and non-contiguous (right) ICs identified by SCA of TRPV1-4 channels. Contiguous ICs are labeled and projected in different colors, whereas non-contiguous ICs are shown in white. (A) Top: SCA results of the full-length TRPV1-4 channel dataset, which identified 6 contiguous ICs, Core IC1, Core IC2, MPD IC, and ARD IC1-3, as well as 15 non-contiguous ICs. Core IC1 and Core IC2 were classified as a single sector, the TRPV1-4 core sector, due to their mutual covariance. Bottom: Average first-order conservation of residues in each of the spatially contiguous independent components (ICs) identified in the SCA of TRPV1-4 channels. Individual ICs are not distinguishable by conservation alone, indicating that covariation drives the observed sector decomposition. Conservation is calculated using the Kullback-Leibler relative entropy (Di) (See methods). Error bars represent one standard deviation. (B) SCA results for the full length TRPV1 channel dataset, which identified contiguous ICs that were homologous to those in TRPV1-4. Core IC1 and Core IC2 of TRPV1 were homologous to those in TRPV1-4, whereas the MPD and ARD ICs of TRPV1 represented the combination of the TRPV1-4 MPD IC with ARDIC3 and ARD IC1 with ARD IC2, respectively. (C) SCA results of the full length TRPV2-4 dataset, which identified the same contiguous ICs as in the TRPV1-4 analysis, indicating that TRPV1 does not drive the observed sector decomposition within this family. (D) IC submatrix (left) and sector projections obtained from the SCA of the transmembrane (TM) domain of the TRPV1 channel dataset. Core IC1-5 exhibited mutual coevolutionary coupling and formed the TRPV1 core sector. The combination of Core IC1 and Core IC3-5 formed a structure homologous to the TRPV1-4 Core IC1, whereas the Core IC2 of the transmembrane TRPV1 SCA was homologous to those of the full length SCAs. The remaining 5 mutually coevolving ICs formed the TRPV1 accessory sector, which does not show homology to previously identified sectors and ICs.

**Figure S3.**
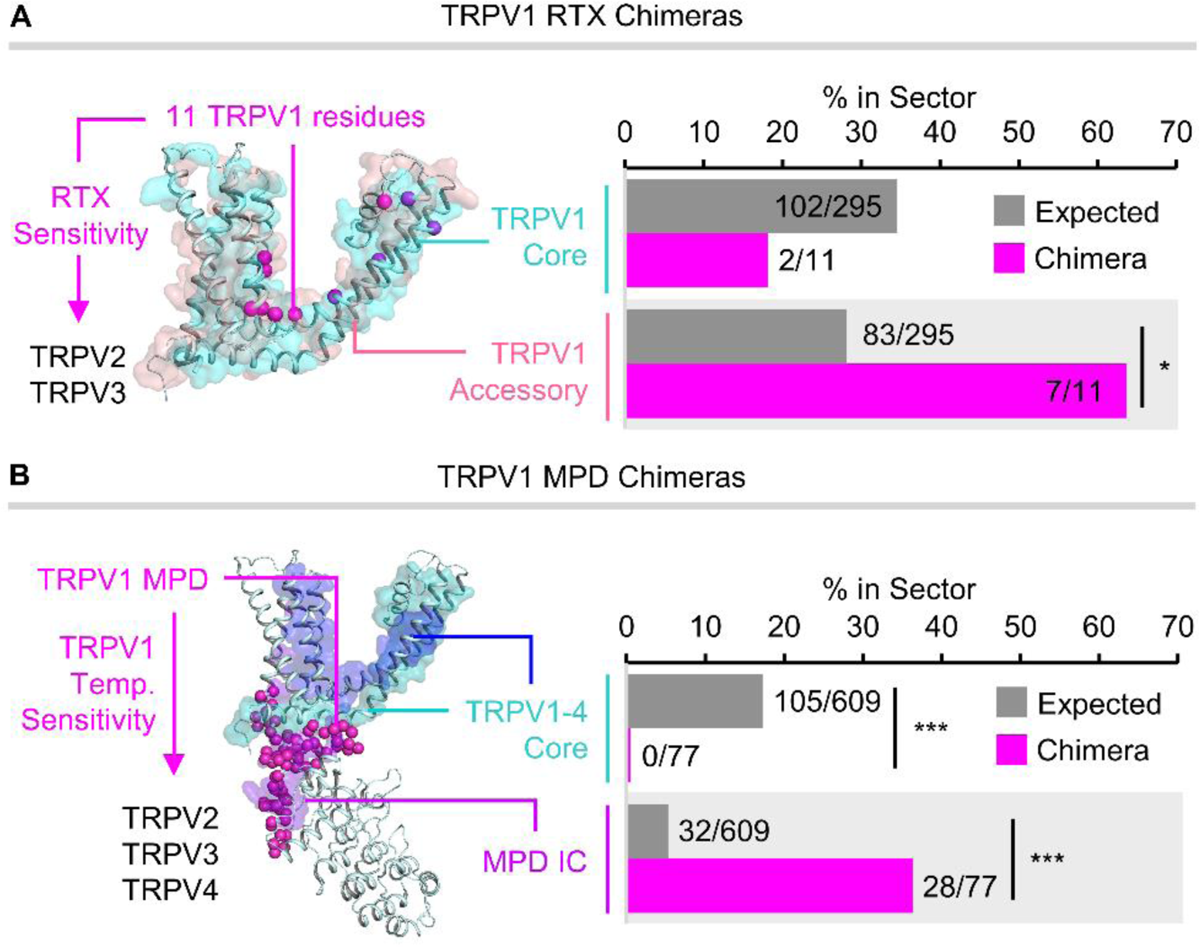
Portable sensory structures identified by chimera experiments map to accessory sectors in TRPV1-4. (A) Structural comparison (left) and percent overlap (right) of the TRPV1 core (cyan) and accessory (pink) sectors with the minimal set of 11 rat TRPV1 residues (Y511, S512, E513, M547, T550, E570, L577, V596, L630, I661, and Y671) (Yang et al., 2016; Zhang et al., 2016, 2019) that are sufficient to introduce RTX sensitivity in chimeras when transplanted to TRPV2 and TRPV3 (magenta spheres). (B) Structural comparison (left) and percent overlap (right) of the TRPV1-4 core sector (Core IC1, cyan; Core IC2, blue) and membrane-proximal domain (MPD) IC with the rat TRPV1 MPD (H358-F434), which was found to confer the activation enthalpy and temperature activation threshold of TRPV1 when transferred to chimeras in TRPV2-4 backgrounds (Yao et al., 2011). All projections are performed on the reference structure of rat TRPV1 (PDB ID: 3j5p, Cao et al., 2013). Expected values represent the percent overlap with the sector expected by random chance, which corresponds to the percent of the protein occupied by the sector (*p < 0.05, **p < 0.01, ***p <0.001, Fisher’s exact test).

**Figure S4.**
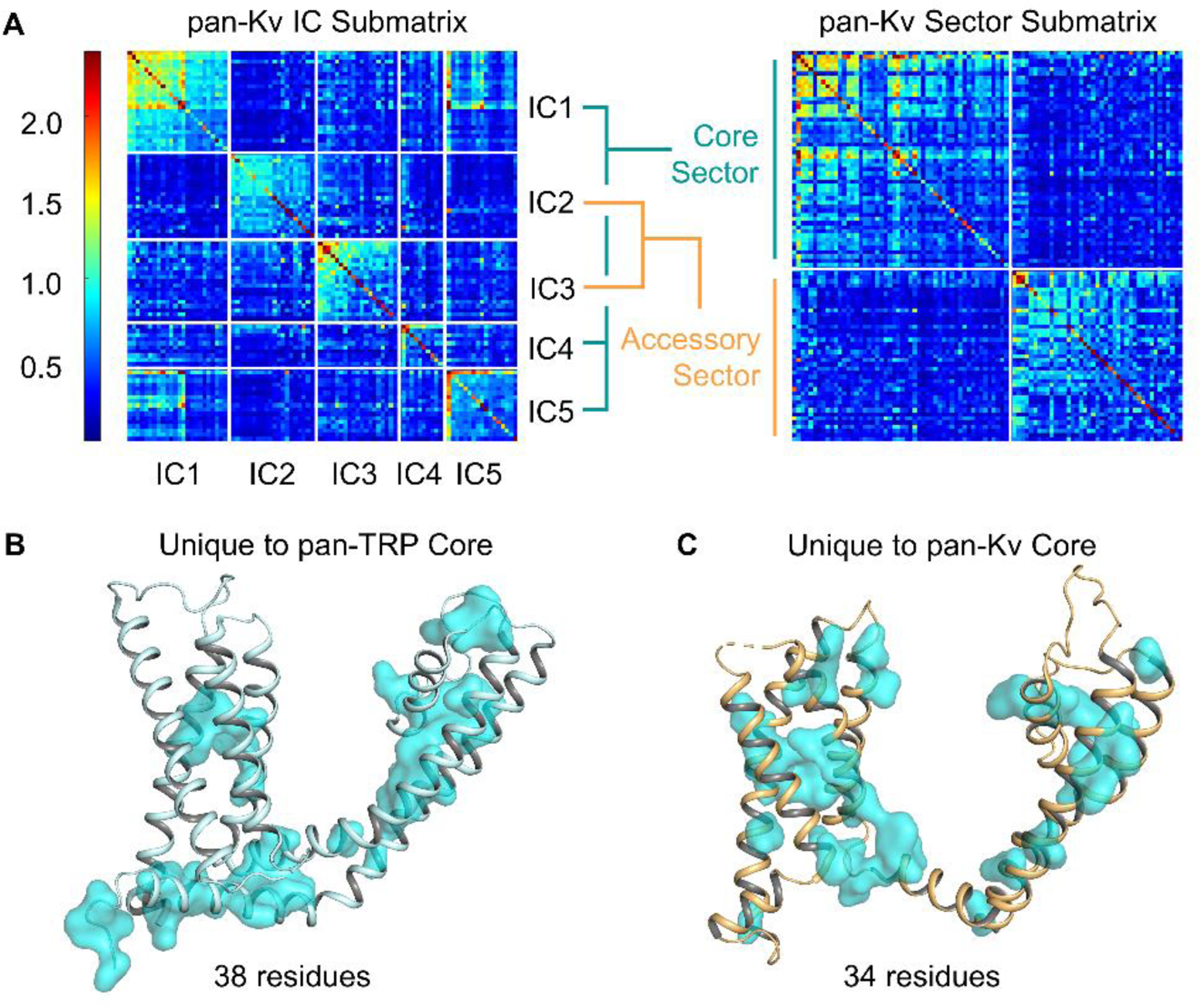
SCA of Kv channels and comparison of the pan-Kv and pan-TRP core sectors. (A) The IC submatrix (left) of the SCA of domain swapped Kv channels showed five ICs which were grouped into the pan-Kv core sector (IC1, IC4, and IC5) and the pan-Kv accessory (IC2 and IC3) based on their patterns of mutual coevolutionary coupling. When these ICs were grouped into sectors and replotted (right), positions in the core and accessory sectors showed strong intra-sector correlations (on-diagonal boxes) but weak inter-sector correlations (off-diagonal boxes), indicating that the two sectors exhibit mutual coevolutionary independence. (B-D) Comparison of the non-positionally homologous residues in the pan-TRP and pan-Kv core sectors as determined by sequence alignment. (B) Positions unique to the pan-TRP core sector (cyan surfaces) map to the TRP helix and extracellular pore domain when projected onto the structure of rat TRPV1 (PDB ID: 3j5p, Cao et al., 2013). (C) Positions unique to the pan-Kv core sector (cyan surfaces) map to the extracellular pore domain and VSD when projected onto the structure of human Kv7.1 (PDB ID: 6uzz, Sun and MacKinnon, 2020). (D) Covariation among pan-Kv accessory sector residues grouped by localization to the VSD or pore domains. The T(V/I)GYG motif of the selectivity filter (T312-G316 in human Kv7.1) coevolves with a subset of VSD residues involved in voltage sensing, including positively charged residues in S4 (R228, R231, Q234, R237, H240, and R243) (Bezanilla, 2000); positions in the charge transfer center (F167, E170, and D202N) (Lacroix et al., 2014; Tao et al., 2010); and E160 in S2 (Cui, 2016).

**Figure S5.**
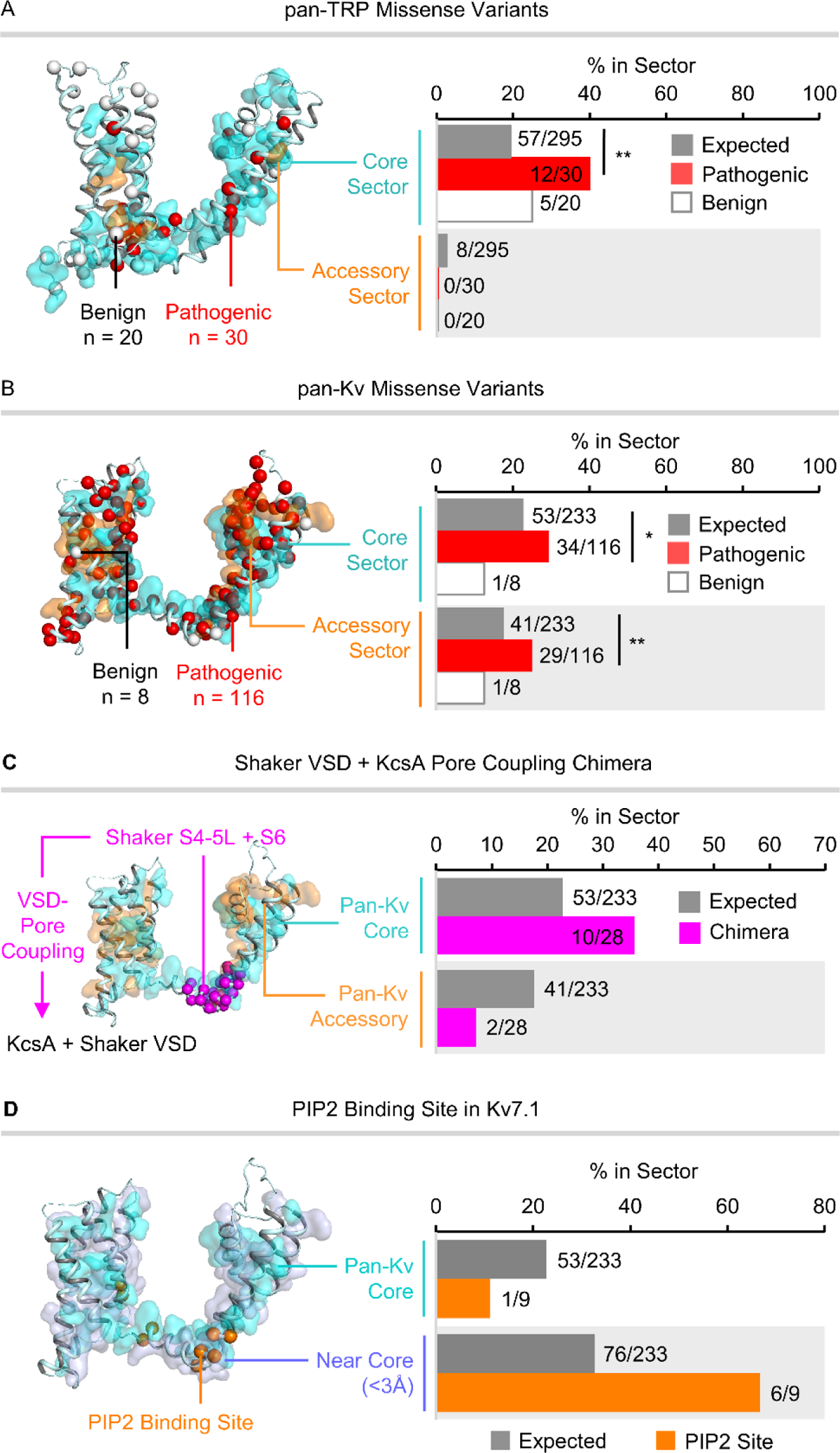
Pathogenic substitutions and coupling interactions map to TRP and Kv sectors. (A) Structural comparison (left) and percent overlap (right) of the pan-TRP core (cyan) and accessory (orange) sectors with the 30 pathogenic and 20 benign missense variants sites reported in the ClinVar database for all human TRP channels. The pan-TRP core sector are enriched in pathogenic but not benign variants. (B Structural comparison (left) and percent overlap (right) of the pan-Kv core (cyan) and accessory (orange) sectors with the 116 pathogenic and 8 benign missense variants sites reported in the ClinVar database (Landrum et al., 2018) for all human domain-swapped Kv channels (Kv1-9). The pan-Kv core and accessory sectors are each enriched in pathogenic but not benign variants. (C) Structural comparison (left) and percent overlap (right) of the pan-Kv core (cyan) and accessory (orange) sectors with short segments in the S4-5 linker and S6 of Shaker (L385-L399 and P473-Y485 in Kv7.1, respectively) that are jointly required to couple the Shaker VSD to the two-transmembrane KcsA channel in chimeras (Lu et al., 2002). The S4-5 linker and S6 segments are slightly but not significantly enriched in pan-Kv core sector residues, suggesting that contact between the pore and core sector is required for coupling. (D) Structural comparison (left) and percent overlap (right) of the pan-Kv core sector (cyan) and positions within 3 Å of the pan-Kv core sector (light blue) with PIP2 binding site residues (orange spheres; F127, F130, G246, T247, R259, Q260, I263, T264, K354, and K358) (Liu et al., 2020) in human Kv7.1. F127 and K358 (not shown) lacked sufficient conservation to be analyzed by SCA and were therefore excluded from the analysis. Rat TRPV1 (PDB ID: 3j5p, Cao et al., 2013) and human Kv7.1 (PDB ID: 6uzz, Sun and MacKinnon, 2020) are used as the reference structures for all TRP and Kv channel projections, respectively. Expected values represent the percent overlap with the sector expected by random chance, which corresponds to the percent of the protein occupied by the sector (*p < 0.05, **p < 0.01, ***p < 0.001, Fisher’s exact test).

**Figure S6.**
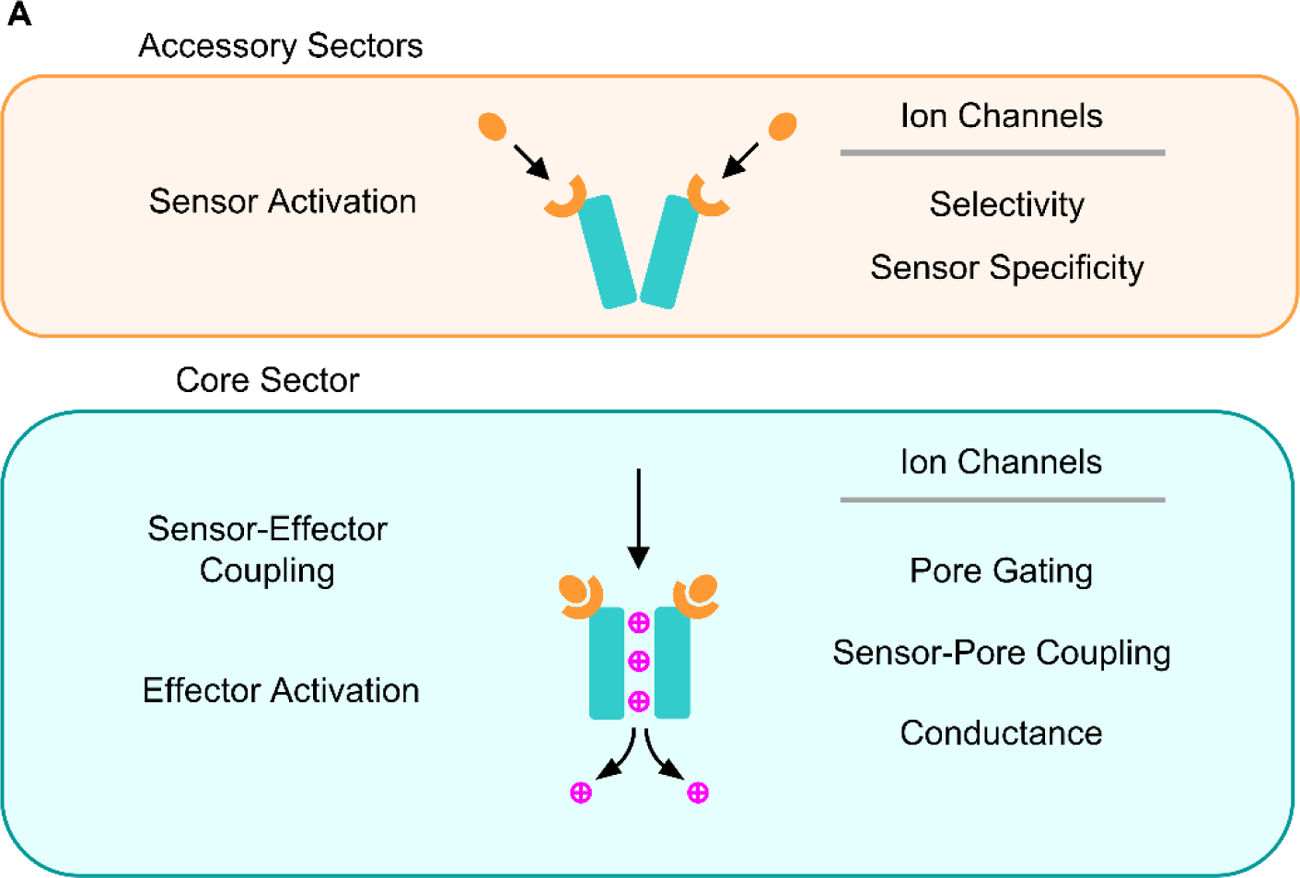
Functional differentiation of core and accessory sectors. (A) Diagram illustrating the functional differentiation of the core and accessory sectors. Receptor-specific sensory functions, including sensor activation and ion selectivity in ion channels, are carried out by the accessory sectors. In contrast, conserved effector functions, including sensor-pore coupling and pore gating in ion channels, are mediated by the core sector.

**Table S1.** Distribution of sequences included in the pan-TRP SCA by species. A total of 1276 TRP channel sequences were comprehensively collected from 56 organisms using BLAST searches against human or *Drosophila melanogaster* TRP channel reference sequences. Species codes correspond to the two- or three-letter code used to annotate sequences in the alignment file.

| Organism | Code | Number of Sequences |
| --- | --- | --- |
| *Homo sapiens* | h | 28 |
| *Mus musculus* | m | 28 |
| *Drosophila melanogaster* | dm | 12 |
| *Caenorhabditis elegans* | ce | 16 |
| *Danio rerio* | z | 22 |
| *Ciona Intestinalis* | ci | 17 |
| *Orbicella faveolata* | of | 37 |
| *Nematostella vectensis* | nv | 31 |
| *Lingula anatina* | lana | 22 |
| *Gallus gallus* | gg | 24 |
| *Branchiostoma belcheri* | bb | 19 |
| *Terrapene carolina triunguis* | tc | 25 |
| *Takifugu rubripes* | tr | 23 |
| *Callorhinchus milii* | cm | 24 |
| *Strongylocentrotus purpuratus* | spur | 39 |
| *Acanthaster planci* | ap | 26 |
| *Penaeus vannamei* | pv | 18 |
| *Limulus polyphemus* | lp | 27 |
| *Aedes aegypti* | ae | 12 |
| *Biomphalaria glabrata* | bg | 15 |
| *Crassostrea virginica* | cv | 36 |
| *Oryctolagus cuniculus* | oc | 25 |
| *Bos taurus* | bt | 26 |
| *Xenopus tropicalis* | xt | 25 |
| *Rattus norvegicus* | rn | 27 |
| *Aptenodytes forsteri* | af | 23 |
| *Loxodonta africana* | lafr | 25 |
| *Exaiptasia pallida* | ep | 14 |
| *Hydra vulgaris* | hv | 17 |
| *Actinia tenebrosa* | at | 21 |
| *Acropora millepora* | am | 35 |
| *Porcillopora damicornis* | pd | 34 |
| *Stylophora pistillata* | spis | 41 |
| *Centruroides sculpturatus* | cs | 19 |
| *Bombus impatiens* | bi | 10 |
| *Monomorium pharaonis* | mp | 10 |
| *Onthophagus taurus* | ot | 14 |
| *Saccoglossus kowalevskii* | sk | 26 |
| *Pomacea canaliculata* | pcan | 18 |
| *Mizuhopecten yessoensis* | my | 25 |
| *Aplysia californica* | acal | 17 |
| *Strongyloides ratti* | sr | 11 |
| *Trichinella spiralis* | ts | 3 |
| *Python bivittatus* | pb | 24 |
| *Anolis carolinensis* | acar | 23 |
| *Equus caballus* | ec | 28 |
| *Panthera tigris* | pt | 19 |
| *Salmo salar* | ss | 36 |
| *Camelus bactrianus* | cb | 26 |
| *Pteropus alecto* | pa | 21 |
| *Physeter catodon* | pcat | 21 |
| *Rhincodon typus* | rt | 18 |
| *Cyprinus carpio* | cc | 28 |
| *Parasteatoda tepidariorum* | pt | 19 |
| *Anneissia japonica* | aj | 22 |
| *Asterias rubens* | ar | 23 |

**Table S2.**
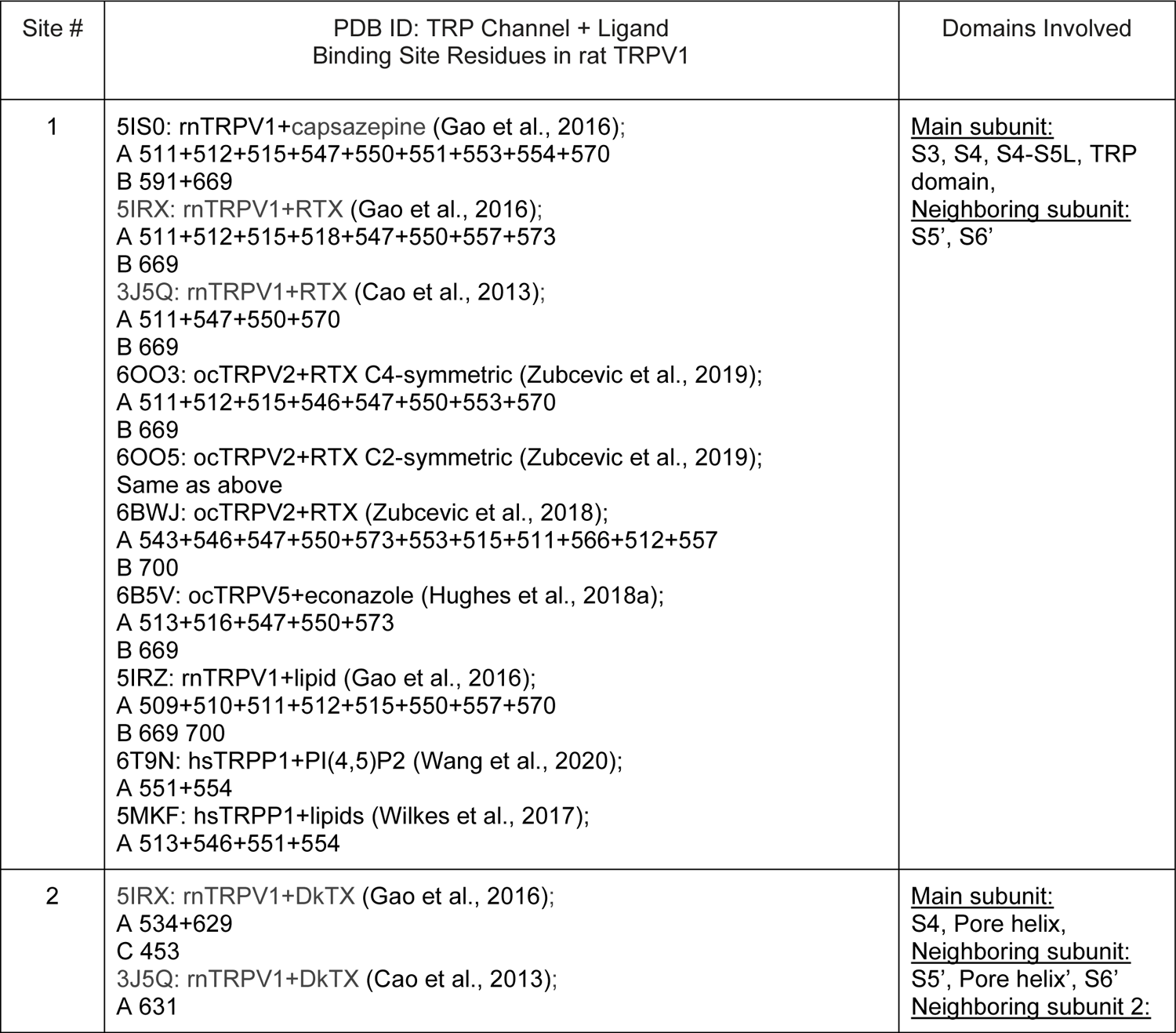
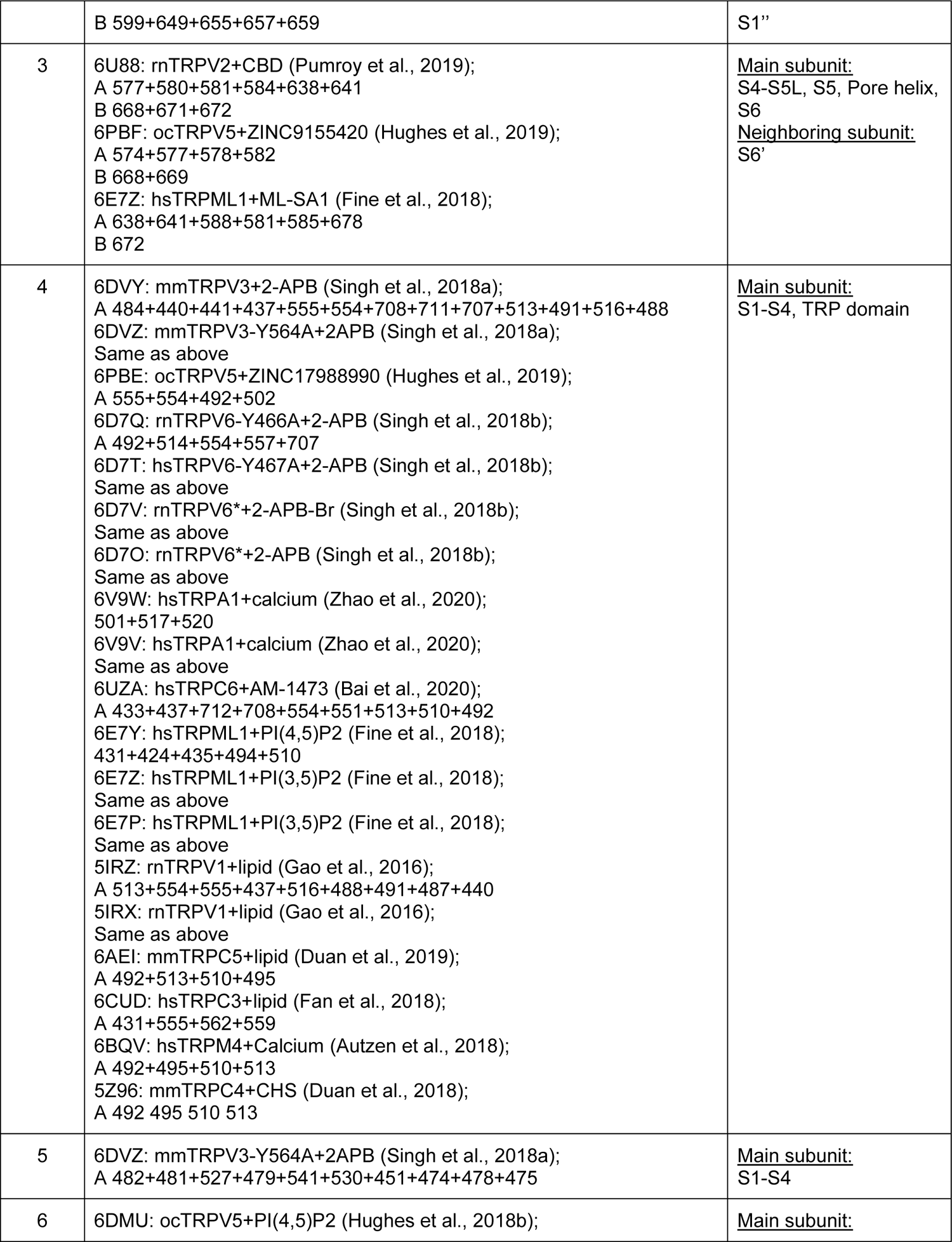
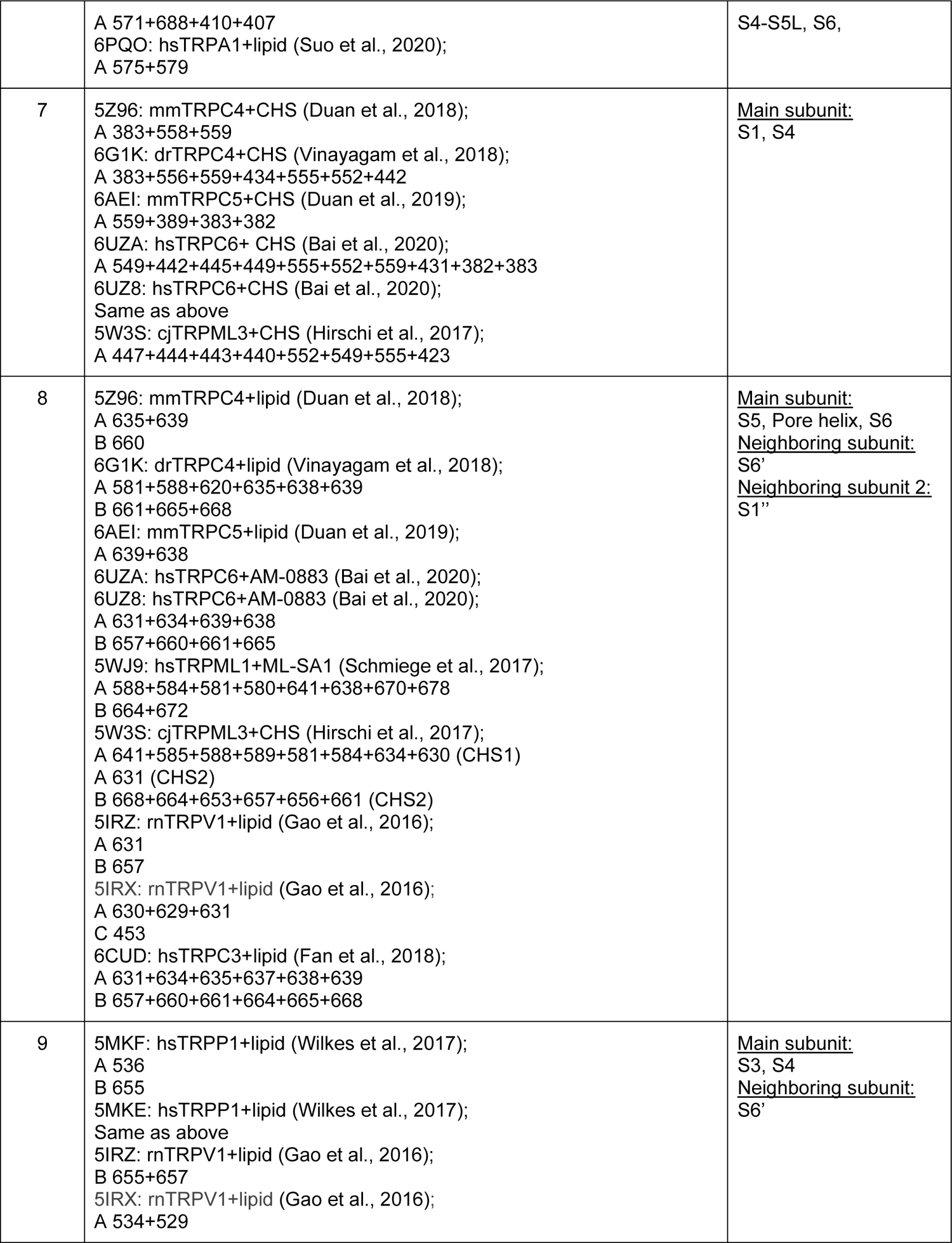
Ligand binding site residues in TRP channels. Binding site residues were determined for 57 ligands in 36 cryo-EM structures of ligand-bound TRP channels using LigPlot2.2+ (Laskowski and Swindells, 2011). Residues making direct contacts with ligands were organized into 9 sites based on structural alignment of the transmembrane domain. Binding site residues were mapped to their corresponding positions in rat TRPV1 and listed under the PDB ID, TRP channel, and ligand associated with each analyzed structure. Chains harboring each residue in the structure (chain A, B, or C) are indicated. For each binding site, the structural domains involved in forming ligand-protein contacts are summarized.

**Table S3.** Input parameters of statistical coupling analysis (SCA). The effective sequence number (M’) represents the number of sequence clusters sharing greater than 80% identity. Maximum fraction of gaps for sequences and positions defined the threshold for including aligned positions or sequences in the final processed alignment. The minimum and maximum identities to the reference sequence (rat TRPV1 for TRP channel SCAs and Kv7.1 for the pan-Kv SCA) were used to eliminate highly dissimilar sequences and calculate sequence weights, respectively. Confidence thresholds indicate the confidence level for determining the top contributing residues in the independent components (ICs) identified by SCA. Non-default parameter values are shown in bold.

| SCA | Total Sequences (M) | Effective Sequence Number ($M'$) | Maximum Fraction of Gaps | | Identity to Reference Sequence | | Confidence Threshold |
| --- | --- | --- | --- | --- | --- | --- | --- |
|  |  |  | Positions | Sequences | Minimum | Maximum |  |
| pan-TRP | 1276 | 617 | 0.2 | 0.2 | **0.1** | **0.85** | **0.97** |
| TRPV1-4 | 2029 | 130 | 0.2 | 0.2 | 0.2 | 0.8 | 0.95 |
| TRPV2-4 | 1315 | 55 | 0.2 | 0.2 | 0.2 | 0.8 | 0.95 |
| TRPV1 | 621 | 61 | 0.2 | 0.2 | 0.2 | 0.8 | 0.95 |
| pan-Kv | 1303 | 180 | 0.2 | 0.2 | 0.2 | 0.8 | 0.95 |

